# Alternative somatic and germline gene-regulatory strategies during starvation-induced developmental arrest

**DOI:** 10.1101/2021.04.30.441514

**Authors:** Amy K. Webster, Rojin Chitrakar, L. Ryan Baugh

## Abstract

Nutrient availability governs growth and quiescence, and many animals arrest development when starved. Somatic and germline cells have distinct functions and constraints, suggesting different regulatory mechanisms contribute to an integrated starvation response. Using *C. elegans* L1 arrest as a model, we show that gene expression changes deep into starvation. Surprisingly, relative expression of germline-enriched genes increases for days. We conditionally degraded the large subunit of RNA Polymerase II and found that early but not late somatic transcription is required for survival and that germline transcription does not affect survival. Expression analysis revealed that thousands of genes are continuously transcribed in the soma, though their absolute abundance declines, while the germ line is transcriptionally quiescent with extreme transcript stability. This work reveals alternative somatic and germline gene-regulatory strategies during starvation, with the soma relying on a robust transcriptional response to support immediate survival while the germ line relies on transcriptional quiescence and mRNA stability to maintain future reproductive success.

## INTRODUCTION

Development requires favorable environmental conditions, and diverse animals enter a state of developmental arrest in response to unfavorable conditions (MacRae 2010). Starvation causes cellular quiescence in cells ranging from yeast to human, and some animals arrest development in response to inadequate nutrition (Su *et al.* 1996; Yao 2014). *C. elegans* nematodes hatch as L1 larvae, and in the absence of food they arrest development, providing a valuable model of starvation resistance and developmental arrest. (Baugh 2013). L1 arrest (or L1 diapause) has garnered attention because time spent in arrest does not shorten lifespan upon recovery, as if it is an “ageless” state (Johnson *et al.* 1984). However, worms in L1 arrest actually exhibit signs of aging, but most are reversible upon recovery (Roux *et al.* 2016a). Nonetheless, extended L1 starvation impacts many life-history traits, including brood size and growth rate (Jobson *et al.* 2015), and some effects persist across generations (Jobson *et al.* 2015; Webster *et al.* 2018). These observations suggest that starvation takes a toll on both somatic and germline cells. These cells have different metabolic demands, developmental constraints, and organismal functions, but how their starvation responses are tailored to their physiology is unknown.

L1 arrest is accompanied by changes in transcriptional regulation and gene expression. Gene expression profiles of mRNA change rapidly early in L1 arrest (within hours), as the starvation response is mounted (Baugh *et al.* 2009). A number of transcriptional regulators, including transcription factors, are required to support starvation survival, suggesting a critical role of transcriptional regulation (Baugh and Sternberg 2006; Zhong *et al.* 2010; Fukuyama *et al.* 2012; Baugh 2013; Cui *et al.* 2013; O’rourke and Ruvkun 2013; Kaplan *et al.* 2015; Murphy *et al.* 2019; Baugh and Hu 2020). However, gene expression dynamics that occur beyond 24 hours of starvation, and time-of-action of transcriptional regulation for supporting survival, are largely unknown.

Transcriptional regulation differs between somatic and germline tissues. In *C. elegans, z*ygotic mRNA transcription begins in somatic cells at the two- to four-cell stage of embryogenesis (Seydoux and Fire 1994; Baugh *et al.* 2003), whereas transcription is repressed by PIE-1 in the P lineage that gives rise to primordial germ cells (PGCs) (Mello *et al.* 1996; Seydoux *et al.* 1996; Seydoux and Dunn 1997). Global repression of transcription in the early embryonic germ line is conserved among metazoa, apparently preventing specification of somatic fates (Wang and Seydoux 2013). Some zygotic mRNA transcription occurs in PGCs during mid-embryogenesis in *C. elegans*, but PGCs remain largely transcriptionally quiescent compared to somatic cells (Wang and Seydoux 2013). Upon hatching, L1 larvae have 558 cells, two of which are the PGCs, Z2 and Z3, that give rise to the germ line. DAF-16/FoxO, DAF-18/PTEN, and the AMP-activated kinase AMPK are each required to arrest somatic cell divisions during L1 starvation, but DAF-18 and AMPK are also required to arrest germ-cell divisions though DAF-16 is not (Baugh and Sternberg 2006; Fukuyama *et al.* 2006; Fukuyama *et al.* 2012). Likewise, DAF-18 and AMPK are required for transcriptional repression of germ cells in arrested L1s (Demoinet *et al.* 2017; Fry *et al.* 2020). Pol II accumulates at promoters of growth genes during L1 arrest (Baugh *et al.* 2009), where it is poised to initiate transcription in response to feeding (Maxwell *et al.* 2014), but the anatomical site of action is unknown. In both somatic and germline cells, transcription increases dramatically upon feeding, supporting post-embryonic development (Furuhashi *et al.* 2010; Maxwell *et al.* 2012; Butuci *et al.* 2015; Fry *et al.* 2020). The extent to which somatic and germline gene regulation share common or distinct mechanisms and regulatory strategies during developmental arrest is unclear.

Here we performed mRNA-seq on starved L1 larvae over twelve timepoints spanning the entirety of L1 arrest. We found that gene expression continues changing throughout starvation, with a rapid initial response followed by a long relatively slow response. Target analysis suggests that known transcriptional regulators mostly act early in starvation, but large numbers of germ line-expressed genes are relatively up-regulated late in arrest. We used selective degradation of AMA-1/Pol II in the soma and germ line to show that transcription during the first two days of starvation in the soma, but at no point in the germ line, is required to support starvation survival. This is consistent with a critical role of transcription in mounting the starvation response. Late in starvation, transcription of thousands of genes continues in the soma as a way of maintaining the initial starvation response while the overall amount of mRNA per animal declines. In contrast, the germ line is transcriptionally inactive throughout arrest, and the relative increase in expression of germline-enriched genes is driven by transcript stability. Collectively, our results suggest that the soma and germ line use distinct regulatory strategies to support organismal fitness during starvation-induced developmental arrest.

## RESULTS

We performed mRNA-seq on whole, starved L1 larvae over time to determine gene expression dynamics throughout L1 arrest. The time series started approximately two hours prior to hatching (−2 hr) to capture the onset of starvation and extended 12 days beyond that (Fig 1A). Because expression dynamics slow within 12 hours (Baugh *et al.* 2009), we collected time points densely early in arrest and sparsely late in arrest (Fig 1A). ~50% of larvae hatched between zero and two hours, and ~80% were hatched by four hours (Fig 1B), reflecting consistent synchrony and staging. About 80% of larvae were still alive at eight days, but survival dropped to about 30% at 12 days (Fig 1C). Thus, the time series spans the entirety of L1 arrest, from hatch to death.

**Figure 1:**
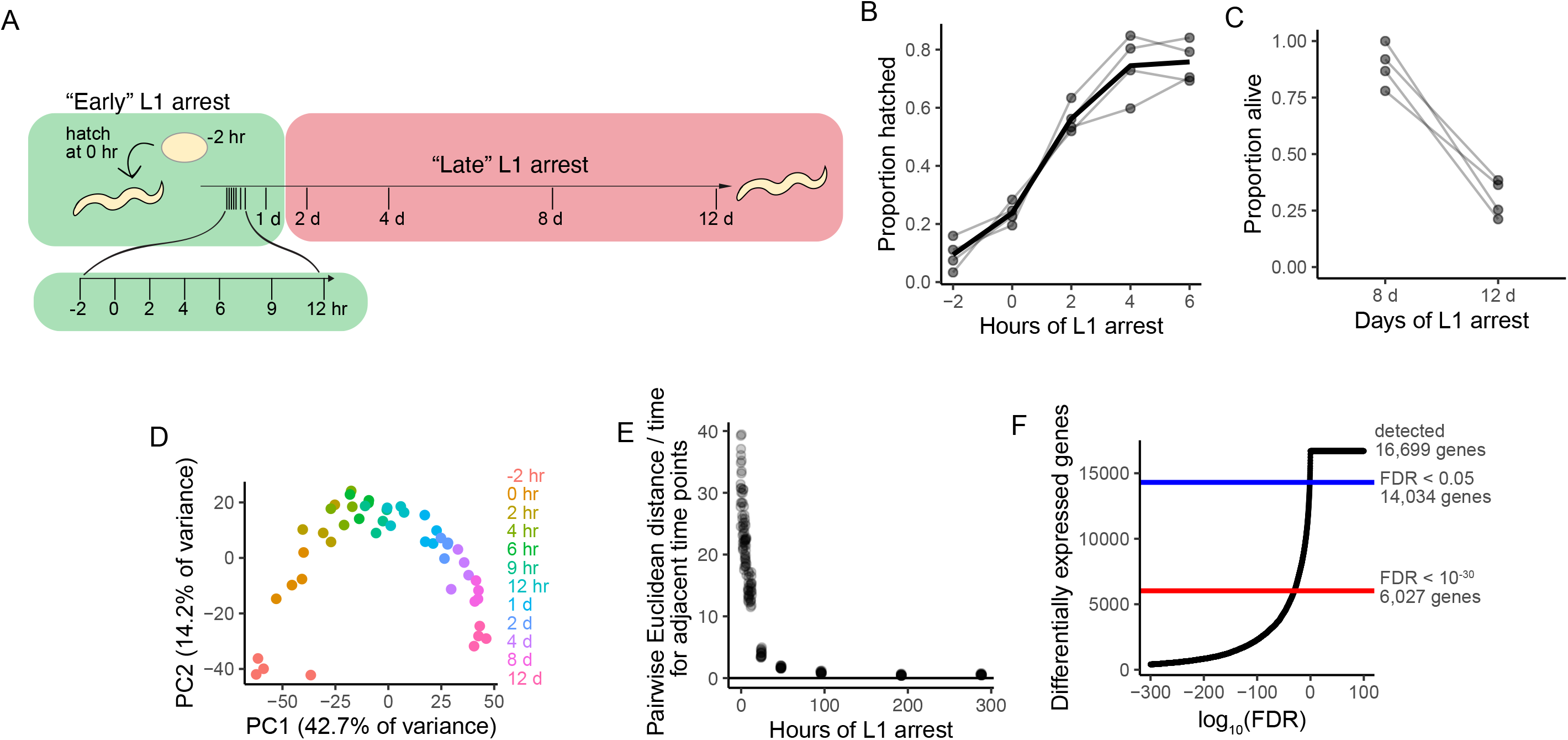
Gene expression changes deep into starvation. A. Time points collected for mRNA-seq time series. B. Hatching efficiency of all replicates at early time points. C. Survival of all replicates at late time points. D. Principal Component Analysis of RNA-seq data for all time points and replicates. E. Rate of change of gene expression based on Euclidean distance. F. Differentially expressed genes as a function of the false-discovery rate.

### Gene expression dynamics throughout starvation

We performed principal component analysis (PCA) as an initial evaluation of expression dynamics. Biological replicates clustered together, as expected, and time points were ordered based on the duration of L1 arrest (Fig. 1D). A rapid response to starvation was evident in the early hours after hatching, as expected (Baugh *et al.* 2009). However, time points including day 1 and beyond were clearly distinct from earlier time points, revealing that gene expression continues changing late in starvation. We measured the rate of change between adjacent time points and found that it decreases dramatically during starvation, with a major inflection near 24 hours (Fig 1E). These results suggest a rapid early response to starvation followed by a much slower late response extending until death.

Strikingly, the vast majority of genes are differentially expressed during starvation. Over 84% of detected genes (14,034 genes) were differentially expressed at a false-discovery rate (FDR) of 0.05, and over 35% (6,027 genes) were differentially expressed at a highly stringent cutoff of 10^−30^ (Fig 1F, Supplementary File 1). These results demonstrate the profound effect of starvation on gene expression.

Despite pervasive effects of starvation, cluster analysis revealed relatively simple temporal patterns. The clustering algorithm used produced 129 clusters for the 6,027 most significantly affected genes, but many of them are distinguished by relatively minor differences in timing or include only a few genes (Supp Fig 1). The ten largest clusters include about two-thirds of genes, and they show the predominant expression patterns present in the full dataset, which broadly consist of genes monotonically increasing or decreasing in relative expression (Fig 2A). The most complex common pattern is an increase followed by a decrease with a single peak early in starvation (*e.g.*, clusters 5, 9, and 10), and other more complex patterns are either very rare or absent (Fig 2B; Supp Fig 1). These observations suggest the gene-regulatory network controlling the starvation response is relatively shallow compared to developmental regulatory networks.

**Figure 2:**
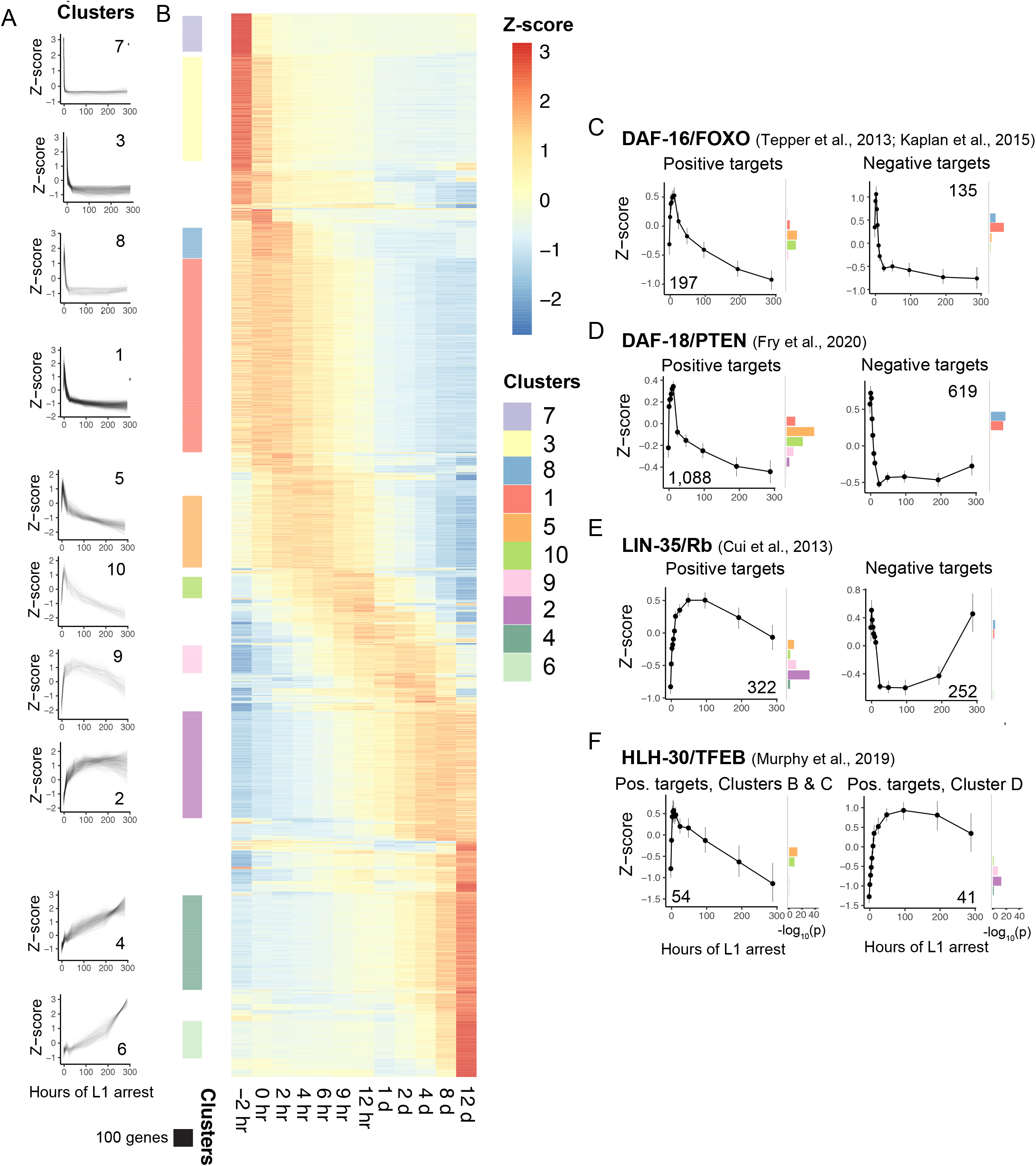
Expression analysis reveals temporal patterns of regulatory activity. A. Z-score-normalized expression dynamics over time for the ten largest clusters. B. Heatmap of all clustered genes (FDR < 10^−30^), sorted by cluster similarity and color-coded by z-score. Colored bars to the left correspond to genes in clusters shown in A. C-F. Gene expression dynamics for known targets of important transcriptional regulators. Number of genes included is inset on each graph. Lines indicate the mean z-score and 99% confidence intervals for all genes in each group. To the right of each graph, -log_10_ enrichment p-values are plotted for the top ten clusters.

### Known transcriptional regulators mostly act early in starvation

Previous studies have identified regulators of L1 starvation survival, including transcription factors and signaling molecules that affect transcription factor activity (Baugh and Sternberg 2006; Zhong *et al.* 2010; Fukuyama *et al.* 2012; Baugh 2013; Cui *et al.* 2013; O’rourke and Ruvkun 2013; Kaplan *et al.* 2015; Murphy *et al.* 2019; Baugh and Hu 2020). We determined expression profiles of known targets (direct and indirect) of critical regulators to shed light on when they are most active. We focused our analysis on DAF-16/FoxO, DAF-18/PTEN, LIN-35/Rb, and HLH-30/TFEB because loss of each severely compromises starvation survival and genome-wide expression data for each mutant in L1 arrest is available (Cui *et al.* 2013; Tepper *et al.* 2013; Kaplan *et al.* 2015; Murphy *et al.* 2019; Fry *et al.* 2020). Positively regulated targets were expressed at their highest levels at different times after hatching, with DAF-16, DAF-18, and some HLH-30 targets peaking between six and 12 hours of arrest, and LIN-35 and other HLH-30 targets peaking between two and four days of L1 arrest (Fig 2C-F), consistent with relatively early and late function, respectively. SKN-1/Nrf and AMPK targets were identified in later developmental stages without starvation (Steinbaugh *et al.* 2015; El-Houjeiri *et al.* 2019), but these targets exhibit peak expression very early in L1 arrest as if these factors help mount the starvation response (Supp Fig 2D-E). In general, positive targets (genes down-regulated in the mutant) peaked after hatching and declined in expression deep into arrest. Negative targets typically declined immediately upon hatching, and in some cases later increased in relative expression deep into arrest (*e.g.*, LIN-35). Like the global dynamics of the starvation response (Fig 1E), these patterns suggest that the transcriptional response to starvation driven by known regulators is largely mounted early. In contrast, regulation accounting for the relative increase in expression observed for hundreds of genes deep in starvation is unknown.

### Differential regulation of germline and somatic genes deep into developmental arrest

Three of our six largest clusters (clusters 2, 4, and 6) were expressed at relatively low levels upon hatching, exhibited peak expression levels beyond four days of starvation, and either maintained or increased expression levels up to 12 days of starvation. These clusters were enriched with genes expressed in several tissues related to reproduction, including “germ line”, “gonad primordium”, and “reproductive system” (Fig 3A) (Angeles-Albores *et al.* 2016), which was surprising given that germ cells are considered relatively transcriptionally quiescent during L1 arrest. Although we assayed whole worms, we further assessed if genes typically expressed in the soma or germ line exhibited distinct expression patterns. We used single-cell RNA-seq data from embryos to define gene sets enriched in somatic or primordial germ cells (PGCs) (Packer *et al.* 2019). PGC-enriched genes were largely expressed at peak levels late in starvation (Fig 3B), and this was robust to defining PGC enrichment with increasing stringency (Supp Fig 3). In agreement, the PGC gene set was overrepresented in seven clusters, including clusters 2, 4, and 6, all of which were stable or increasing over time during starvation. In contrast, soma-enriched genes were over-represented in clusters with peak expression within the first few hours of starvation (clusters 1, 3, 5, and 7). Collectively, these results indicate that PGC-enriched genes tend to increase in relative expression throughout arrest, while soma-enriched genes tend to peak early and decrease.

**Figure 3:**
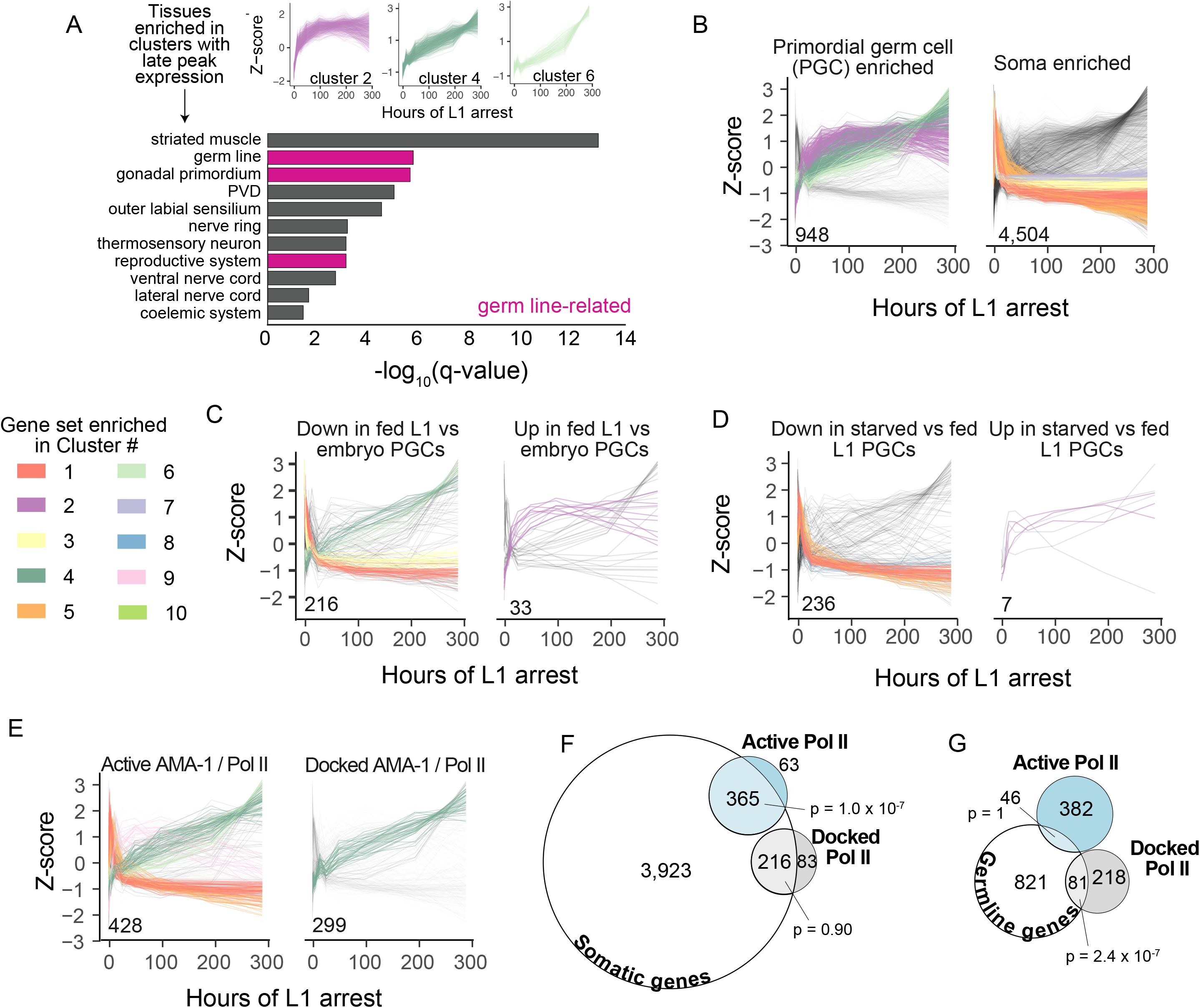
Somatic and germline genes have different patterns of regulation during starvation. A. Tissue enrichments for clusters 2, 4, and 6, which have peak expression late in starvation. Germline-related tissues are highlighted in pink. B-E. Z-scores over time are plotted for all individual genes in the indicated group and clustered dataset. The number of genes is inset on each graph. Genes are color-coded (see legend) by cluster if the cluster is enriched in the gene group with all other genes in grey. B,E. Cluster enrichment color-coded if hypergeometric p < 0.01. C,D. Cluster enrichment color-coded if hypergeometric p < 0.05. F. Venn diagram of germ line-enriched genes plotted in B and genes with active (hypergeometric p = 1) or docked (hypergeometric p = 2.4 x 10^−7^) Pol II. G. Venn diagram of soma-enriched genes plotted in B and genes with active (hypergeometric p = 1.0 x 10^−7^) or docked (hypergeometric p = 0.90) Pol II.

We extended our analysis with a pair of published datasets examining expression in PGCs sorted from L1-stage larvae, including 1) genes differentially expressed between embryonic and fed L1 PGCs and 2) genes differentially expressed between fed and starved L1 PGCs (Lee *et al.* 2017). As expected, genes down-regulated between in L1 compared to embryonic PGCs were over-represented in clusters that decrease in expression within hours of hatching (clusters 1 and 3), though they were also enriched in clusters that increase late (clusters 4 and 6) (Fig 3C). In contrast, the majority of genes up-regulated in L1 relative to embryonic PGCs were part of the late clusters 2, 4 and 6, with cluster 2 significantly over-represented (Fig 3C). Genes down-regulated in starved compared to fed PGCs were over-represented among clusters with peak expression within the first few hours of L1 arrest (clusters 1, 5, and 8), though the late-peaking cluster 6 was also enriched (Fig 3D). In contrast, six of the seven genes that are up-regulated in starved L1 PGCs exhibited peak expression late in arrest, including three genes significantly enriched in cluster 2 (Fig 3D). Together these observations suggest that genes with differential expression in L1 PGCs, due to either developmental regulation or starvation, show similar patterns early in L1 arrest in our whole-animal data. They also further support the conclusion that many germline genes increase in relative expression levels deep into starvation.

Distinct temporal patterns of steady-state expression for germ line and soma-enriched genes could be driven by active transcription or differences in transcript stability. To gain insight on this distinction, we used previously defined gene sets early in L1 starvation in which Pol II is ‘active’ or ‘docked’ (Maxwell *et al.* 2014). Active genes accumulate Pol II in the gene body, have evidence of elongation activity and mRNA expression, and are enriched for starvation-response genes. In contrast, docked genes are ‘poised’ in that they accumulate Pol II just upstream of the transcription start site, have little to no elongation activity or mRNA expression, and tend to be immediately up-regulated during recovery from starvation. Genes with active Pol II early in starvation are enriched with clusters with peak expression early, mid-way, and late in starvation (clusters 1 and 5, 9, and 4 and 6, respectively) (Fig 3E). In contrast, genes with docked Pol II were enriched for the late-peaking cluster 4 alone (Fig 3E). We found significant enrichment between differentially expressed somatic genes and active but not docked Pol II (Fig 3F), consistent with active transcription of somatic genes early in starvation. We also found significant enrichment between differentially expressed germ line-enriched genes and docked but not active Pol II (Fig 3G). These associations suggest soma-enriched genes are actively transcribed and germ cells are transcriptional quiescent early in starvation, as expected, but whether the up-regulation of germline and docked genes late in starvation is driven by active transcription remains unclear.

### Early somatic but not germline transcription supports starvation resistance

To causally ascertain the role of transcription in the soma and germ line throughout starvation, we used the auxin-inducible degron (AID) system to selectively degrade the large subunit of Pol II, AMA-1. We used CRISPR to tag AMA-1 with a degron (Fig 4A), and we combined this allele with somatic (*Peft-3::TIR1)* or germline (*Pgld-1::TIR1)* transgenes to enable auxin-dependent degradation of AMA-1 in those tissues (Zhang *et al.* 2015; Kasimatis *et al.* 2018). There was a dramatic reduction in hatching efficiency when 0.1 mM auxin was added to *Peft-3::TIR1; ama-1::AID* embryos (Fig 4B), consistent with the embryonic arrest caused by transcription inhibition (Edgar *et al.* 1994; Powell-Coffman *et al.* 1996). These embryos hatched normally when only solvent was added (ethanol), and wild-type (N2) and *Peft-3::TIR-1* embryos hatched at high frequency when auxin was added. When auxin was added to *Pgld-1::TIR1; ama-1::AID* embryos, they developed and hatched, consistent with the germ line being dispensable to embryonic development and largely transcriptionally quiescent. Post-embryonic development depends on somatic and germline transcription, and auxin treatment caused larval growth arrest in *Peft-3::TIR1; ama-1::AID* worms and sterility in *Pgld-1::TIR1; ama-1::AID* worms (data not shown), as expected. Together these results suggest the AID system is effective at selectively degrading AMA-1 in the soma or germ line throughout development.

**Figure 4:**
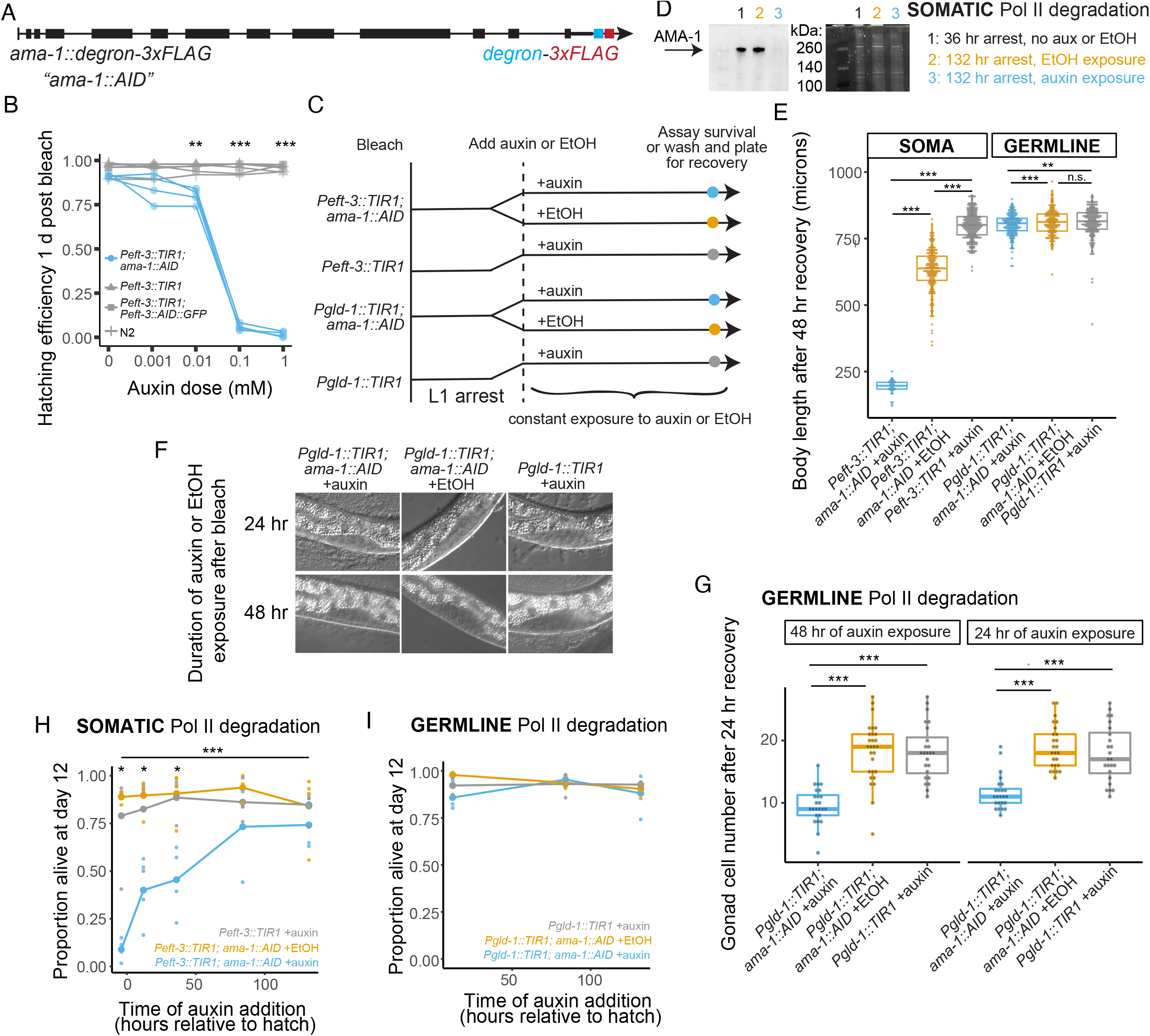
Early somatic transcription supports starvation survival, but germline and late somatic transcription are dispensable. A. Design of *ama-1* degron strain referred to as *ama-1::AID*. The degron and 3x-FLAG sequence were inserted at the C-terminus of the endogenous *ama-1* locus. B. Embryonic hatching efficiency of *Peft-3::TIR1; ama-1::AID* and control strains with different doses of auxin. T-tests between *Peft-3::TIR1; ama-1::AID* (n = 4) and each control strain (n = 2) were performed for each dose. ***p < 0.001, **p < 0.01 compared to each control. C. Experimental design for survival and recovery experiments. Auxin addition to *Peft-3::TIR1; ama-1::AID* degrades AMA-1 in the soma; auxin addition to *Pgld-1::TIR1; ama-1::AID* degrades AMA-1 in the germ line. Ethanol (EtOH) addition and auxin addition to *Peft-3::TIR1* and *Pgld-1::TIR1* are controls. D. Western blot of *Peft-3::TIR1; ama-1::AID* after 36 hours of arrest without auxin exposure, and with and without auxin exposure at 132 hours of arrest. For 132 hours samples, auxin or ethanol exposure began at 36 hr. Total protein for the corresponding part of the gel is shown to the right. The full western blot and total protein gel including two additional biological replicates are shown in Supplementary Figure 4. E. Body length of worms after 48 hours recovery with food following exposure of arrested L1s of the indicated strains to auxin or EtOH for one hour. Linear mixed-effects model with conditions as fixed effect and replicate as random effect. ***p < 0.001, **p < 0.01, n.s. not significant. F. Representative images of the gonadal cells scored in G. G. Number of gonadal cells in larvae of indicated genotypes recovered with food for 24 hours following L1 arrest. Prior to recovery, L1s arrested for 12 hours were exposed to 24 hours of auxin or ethanol, or embryos immediately post-bleach were exposed to 48 hours of auxin or ethanol, both in the absence of food. T-tests, ***p < 0.001. H-I. Survival at 12 days of L1 arrest following constant exposure of indicated genotypes to auxin or ethanol at indicated time point. F. Pairwise t-tests were performed for *Peft-3::TIR1; ama-1::AID* + auxin compared to *Peft-3::TIR1; ama-1::AID* + EtOH and *Peft-3::TIR1* + auxin at each time point. *p < 0.05 in both t-tests. One-way ANOVA across all time points for *Peft-3::TIR1; ama-1::AID* + auxin, ***p < 0.001.

The AID system also effectively eliminates AMA-1 during L1 arrest (Fig 4C), affecting subsequent somatic and germline development. AMA-1 was efficiently degraded in starved L1 larvae when auxin was added to *Peft-3::TIR1; ama-1::AID* (Fig 4D and Supp Fig 4A-B), consistent with the AID system in other developmental stages (Zhang *et al.* 2015), and suggesting a genetic null for *ama-1*. Furthermore, degradation of AMA-1 in the soma during L1 arrest subsequently prevented larval growth during 48 hours on food in the absence of auxin (Fig 4E), consistent with somatic transcription being essential to development. Notably, *Peft-3::TIR1; ama-1::AID* worms grew slower in the absence of auxin than *Peft-3::TIR1* worms with auxin. This is consistent with some auxin-independent background degradation of AMA-1, an effect that has been documented for other degron-tagged proteins in the presence of TIR1 (Schiksnis *et al.* 2020). There was a small but significant effect on body length when auxin was added to *Pgld-1::TIR1; ama-1::AID* worms during L1 arrest, washed thoroughly, and allowed to feed for 48 hours (Fig 4E), suggesting germline development was reduced by AMA-1 degradation in the germ line during L1 arrest resulting in smaller worms. To directly address this, we treated *Pgld-1::TIR1; ama-1::AID* embryos and arrested L1s with auxin, incubated them for 48 or 24 hours, respectively, washed them thoroughly, recovered them with food for 24 hours, and counted the number of gonadal cells (germline and somatic) per worm (Fig 4F-G). Germline degradation of AMA-1 during L1 arrest significantly reduced the number of gonadal cells upon recovery to approximately 10-12 per animal on average. Notably, there are approximately 12 somatic gonad cells at the late L2 stage (Hubbard and Greenstein 2005), suggesting essentially complete inhibition of germ-cell proliferation. Together these results show that the AID system is effective at selectively degrading AMA-1 in the soma or germ line during L1 arrest, and they reveal that doing so inhibits subsequent somatic and germline development, respectively, in fed larvae.

We determined when somatic transcription is essential for starvation survival. We degraded somatic AMA-1 beginning approximately four hours prior to hatching (−4 hours, allowing embryos to hatch and be exposed to auxin for all of L1 arrest), along with 12, 36, 84, and 132 hours after hatching without food, and we scored survival on day 12 (~276 hours). We chose day 12 because starved worms survive longer in the presence of ethanol (Castro *et al.* 2012), which is in auxin and control conditions. Degrading AMA-1 in the soma starting at −4 hours severely limited survival (Fig 4H), as expected given early action of critical transcriptional regulators (Fig 2C-F, Supp Fig 2). Preventing somatic transcription after 12 and 36 hours of L1 arrest also reduced survival, but the effect was diminished compared to adding auxin before hatching (−4 hr), highlighting the importance of transcription during the first 12 hours of L1 arrest. Exposure of wild-type (N2) worms to alpha-amanitin to inhibit transcription starting 12 hours after hatching also modestly reduced survival by day 8, corroborating our results in a complementary system (Supplementary Fig 4C). In contrast, when we degraded AMA-1 starting at 84 and 132 hours, we did not detect a significant decrease in survival, suggesting that late transcription does relatively little to support survival. These results demonstrate that somatic transcription is initially required to mount the starvation response but that transcription is largely dispensable after that, consistent with the observed inflection in gene expression dynamics during early L1 arrest (Fig 1E).

We determined if germline transcription supports starvation survival. (Fig 4I). We added auxin to *Peft-3::TIR1; ama-1::AID* worms 12, 84, and 132 hours after hatching without food. Degradation of somatic AMA-1 at 12 hours had a robust effect on survival (Fig 4H), and the later treatments allowed us to determine whether late transcription of germline genes, a possibility suggested by the time series, impacts survival positively or negatively. However, degradation of germline AMA-1 did not affect survival. This suggests that germ line transcription, if it occurs, is dispensable for starvation survival, though it is required for recovery (Fig 4F,G).

### Transcription occurs in the soma but not germ line days into starvation

We determined whether gene expression changes days into starvation are driven by ongoing transcription or differences in transcript stability. We performed an additional mRNA-seq experiment with samples collected after 36 or 132 hours of L1 arrest, with auxin or ethanol added at 36 hours, and *Peft-3::TIR1* and *Pgld-1::TIR1* strains were included as controls (Fig 5A). Worms had not died due to lack of transcription at 132 hours (Supplementary Fig 5A), facilitating collection of RNA. PCA revealed that time spent in arrest explained more variance in the data than any other factor, with clear separation of 36 and 132 hour samples in principal component 1 (Fig 5B). Principal component 2 separated samples with somatic AMA-1 degraded from all others, demonstrating transcription plays a role in shaping late expression dynamics. Notably, we did not see separation of samples with germline AMA-1 degraded, suggesting relatively little transcription in germ cells, consistent with our survival results (Fig 4I). In addition, all controls clustered together, suggesting that addition of auxin to *Peft-3::TIR1* or *Pgld-1::TIR* alone did not substantially affect gene expression. We identified differentially expressed genes for each strain at 132 hours by comparing auxin and ethanol treatments. For *Peft-3::TIR1; ama-1::AID* and *Pgld-1::TIR1; ama-1::AID* strains, differentially expressed genes indicate transcription-dependent gene expression changes in the soma or germ line, respectively. Using an FDR of 0.05, the results corroborated PCA, identifying thousands of genes with increased or decreased transcript abundance when somatic AMA-1 was degraded, zero genes when germline AMA-1 was degraded, and almost no genes in the control strains (Fig 5C, Supplementary File 1). We conclude that somatic transcription plays a substantial role in shaping gene-expression dynamics late in L1 arrest, and that germ cells remain transcriptionally quiescent deep into arrest.

**Figure 5:**
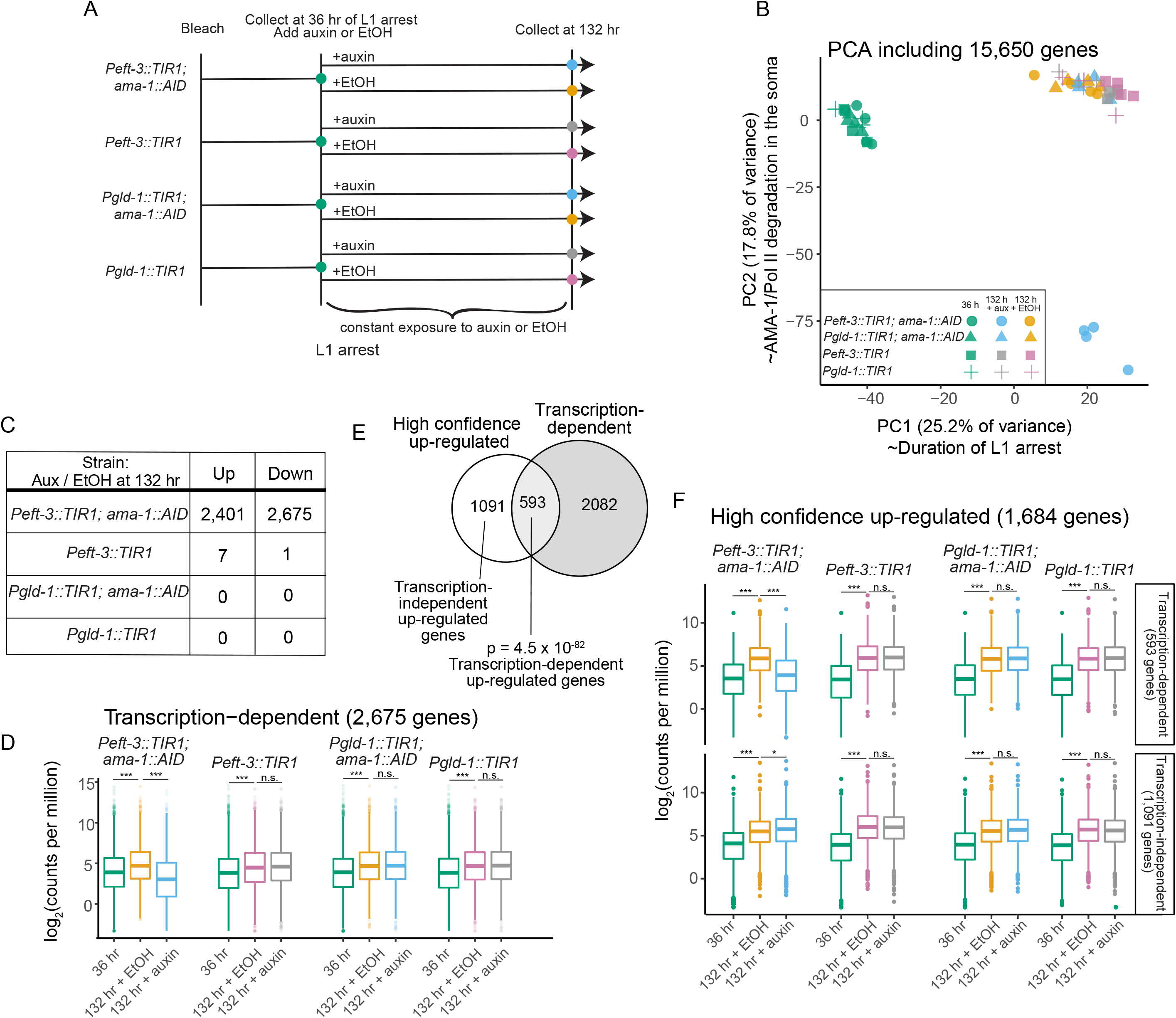
Transcription-dependent gene expression changes occur in the soma but not the germ line late in starvation. A. Experimental design of mRNA-seq experiment. Arrested L1s of indicated genotypes are collected at 36 hours of arrest. After constant exposure to auxin or ethanol from 36 hours onward, arrested L1s of indicated genotypes are collected at 132 hours of arrest. All other conditions are controls. B. Principal component analysis of all conditions and replicates. C. Number of differentially expressed genes for each genotype at 132 hours of L1 arrest depending on whether exposed to auxin or ethanol. D. Log2 counts per million plotted for all transcription-dependent genes (genes down-regulated in *Peft-3::TIR1; ama-1::AID* + auxin compared to *Peft-3::TIR1; ama-1::AID* + EtOH) across all conditions. E. Venn diagram of ‘high confidence up-regulated genes’ (genes up-regulated at 132 hours with EtOH exposure compared to 36 hours in all four genotypes) and ‘Transcription-dependent genes.’ F. Log2 counts per million reads plotted for all ‘High confidence up-regulated genes’, parsed by whether they are transcription-dependent or transcription-independent, as designated in E. E,G. Significance determined using the Kolmogorov-Smirnov test. In each strain background, 36 hours vs 132 hours + EtOH and 132 hours + EtOH vs 132 hours + auxin were compared. *p < 0.05, **p < 0.01, ***p < 0.001.

We were particularly interested in the 2,675 genes with decreased expression when somatic AMA-1 was degraded. On average, these “transcription-dependent” genes were relatively up-regulated between 36 and 132 hours of arrest with ethanol in all four strains (Fig 5D). This temporal increase in expression was abrogated when somatic AMA-1 was degraded, but not when germline AMA-1 was degraded or in control strains. Again, these results demonstrate a role of somatic, but not germline, transcription beyond 36 hours of arrest.

We determined the proportion of genes with increased expression over time due to transcription. 1,684 genes were consistently up-regulated between 36 and 132 hours with ethanol in all four strains, and 35% of these “high confidence up-regulated” genes were also among the 2,675 transcription-dependent genes (Fig 5E). We refer to this overlapping set as “transcription-dependent up-regulated genes.” On average, this gene set behaved like the larger transcription-dependent set (Fig 5D), with an increase in expression over time in all cases but when somatic AMA-1 was degraded, except that the effect sizes were larger for the transcription-dependent up-regulated genes (Fig 5F). These results show that transcription drives increased expression of these genes late in arrest. In contrast, high confidence up-regulated genes that were not also classified as transcription-dependent (“transcription-independent up-regulated genes”) had increased average expression over time even when somatic AMA-1 was degraded, consistent with greater relative transcript stability driving increased relative expression of these genes. We conclude that transcription is the primary cause of increased expression late in starvation for about one-third of up-regulated genes, but that differences in transcript stability drive increases in relative expression for about two-thirds of up-regulated genes.

In our original time series, we found differentially expressed soma-enriched genes exhibit peak expression in the first hours of arrest and are associated with active Pol II, while differentially expressed germ-cell-enriched genes exhibit peak expression days into arrest and are associated with docked Pol II (Fig 3). Strikingly, transcription-dependent genes were most strongly enriched for clusters with peak expression early in arrest (clusters 1, 5, 8, 9, and 10) (Fig 6A). Notably, clusters with increasing expression late in arrest (*e.g.*, clusters 2, 4, and 6) were not enriched, nor were clusters with peak expression in late embryos (clusters 3 and 7). As a subset of transcription-dependent genes, transcription-dependent up-regulated genes were enriched among clusters with peak expression early in arrest (clusters 1 and 8) (Fig 6B). Both gene sets were enriched for somatic genes and Pol II active genes, but not germline genes or Pol II docked genes. Thus, transcription-dependent genes are transcribed late in arrest as a continuation of the early starvation response, even though the majority of them do not increase expression late in arrest. In contrast, transcription-independent up-regulated genes were enriched for cluster 6, which increases in relative expression late in arrest (Fig 6C). Furthermore, they were enriched for germline genes and Pol II docked genes. In addition, the germ line and reproductive system were highly enriched among transcription-independent up-regulated genes (Fig 6D), and the significance of enrichment was substantially greater than for all genes in clusters with late peak expression (Fig 3A). These results support the conclusion that greater relative transcript stability drives late increases among germline-enriched genes rather than transcription.

**Figure 6:**
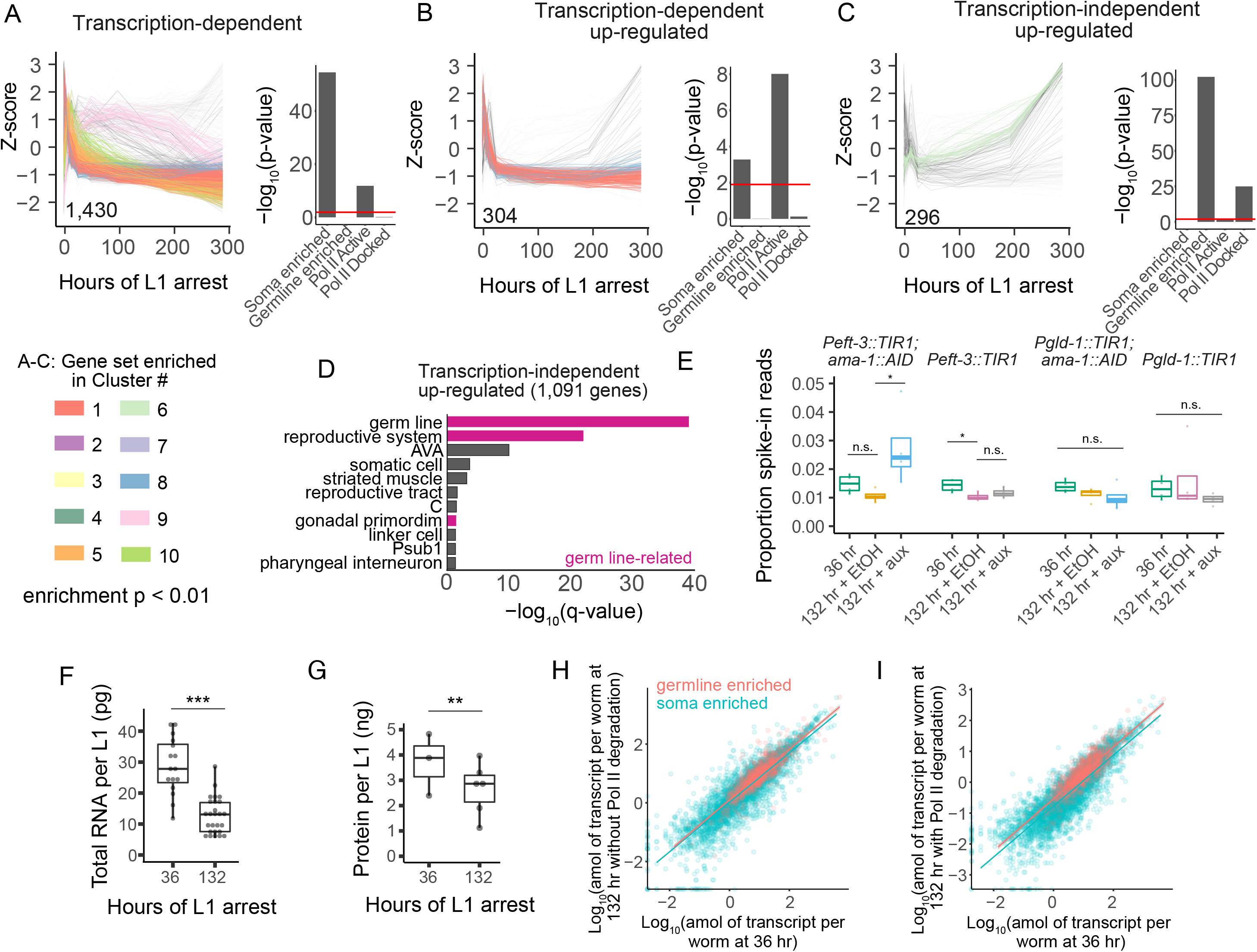
Transcription-independent increased expression late in starvation is driven by relative stability of germline transcripts. A-C. Z-score over time is plotted for all individual genes in the indicated gene group that are part of the clustering dataset. The number of genes plotted is inset on each graph. Genes are color-coded by cluster if that cluster is enriched (hypergeometric p < 0.01) in the gene group. Hypergeometric p-values are plotted for each gene group and its overlap with Soma enriched, Germline enriched, Pol II Active, and Pol II Docked genes (shown in Figure 3). D. Tissue enrichments of all ‘High confidence up-regulated genes’. Germline tissues are highlighted in pink. E. Proportion of reads mapping to spike-in transcripts. One-way ANOVAs across three conditions in each genetic background were performed. If p < 0.05, then pairwise t-tests were performed. *p ≤ 0.05 in t-test. F. Total RNA for all available samples used in RNA-seq separated by duration of starvation with all genotypes and conditions included. G. Total protein for *Peft3::TIR1; ama-1::AID* samples included in western blot. F-G. Linear mixed-effects model with duration of starvation as a fixed effect and biological replicate as a random effect, **p < 0.01, ***p < 0.001. H, I. Log_10_-transformed attomoles of transcript per worm calculated based on spike-ins and total RNA yield per worm for somatic and germline transcripts in the *ama-1::AID;Peft-3::TIR1* strain at 36 hours and 132 hours with ethanol (no degradation of AMA-1) (H) or auxin (degradation of AMA-1) (I). Lines indicate linear regression for each gene group with line width indicating 95% confidence interval.

As a further check that germline-enriched genes are relatively stable compared to soma-enriched genes, we used external standards for absolute quantification of mRNA abundance. During library preparation for the AMA-1 AID mRNA-seq experiment (Fig 5A), we added a pool of 92 synthetic transcripts (spike-ins) at known concentrations based on total RNA content. Because a constant spike-in-to-total RNA ratio was used, determining the proportion of mRNA-seq reads mapping to spike-ins provides a proxy for mRNA content, with a higher proportion mapping to spike-ins indicating a lower mRNA-to-total RNA ratio. *Peft-3::TIR1; ama-1::AID* worms at 132 hours with auxin had a significantly higher proportion of reads mapping to spike-ins compared to the ethanol control, showing that, consistent with expectation, mRNA decreases relative to total RNA when AMA-1 is degraded in the soma (Fig 6E). However, there was not a significant increase in the proportion of spike-in reads between 36 and 132 hours with ethanol in any strain background (Fig 6E), suggesting a relatively constant mRNA-to-total RNA ratio is maintained between 36 and 132 hours. We assessed total RNA levels for samples used in the mRNA-seq experiment. At 132 hours, there was a significant, approximately two-fold decrease in total RNA isolated relative to 36 hours, with RNA isolated from equal numbers of larvae (Fig 6F, Supp Fig 5). These observations together suggest that total RNA declines late in starvation, and that mRNA declines approximately in tandem. In addition to a decline in total RNA, total protein declined between 36 and 132 hours (Fig 6G and Supp Fig 5), and larvae shrink throughout L1 arrest (Hibshman 2017), consistent with a systemic decline in macromolecular content per individual. Finally, using total RNA and spike-ins to normalize mRNA-seq count data to transcripts per worm, we found that absolute expression levels of germline-enriched genes are more resistant to AMA-1 degradation than soma-enriched genes (Fig 6H-I). Both somatic and germline-enriched transcripts declined upon AMA-1 degradation between 36 and 132 hours, but this decline was larger for soma-enriched genes. These results support the conclusion that germline-enriched transcripts are particularly stable, maintaining their relative expression levels deep into starvation despite lack of germline transcription. This is in contrast to soma-enriched genes, which continue to be transcribed as their transcript abundance nonetheless wanes.

## DISCUSSION

We sought to gain deeper insight into the role of transcription in the organismal response to starvation. We used mRNA-seq to characterize gene expression dynamics in whole L1-stage *C. elegans* larvae throughout starvation-induced developmental arrest. We leveraged a variety of genome-scale datasets for integrative analysis of our results, allowing us to infer mechanisms and anatomical sites of regulation. We also took advantage of the AID system to conditionally degrade AMA-1/Pol II in the soma or germ line, assessing the role of transcription in shaping expression dynamics during starvation and its contribution to survival. Together, our results suggest a model (Fig 7) in which early somatic transcription is essential to mount the starvation response in support of survival, and ongoing somatic transcription maintains this response, but late transcription does not appreciably support survival. In contrast to the soma, the germ line remains largely transcriptionally quiescent throughout arrest, but germline transcripts are stable, while somatic transcripts turn over and collectively decline during arrest, leading to a relative increase of germline transcript abundance late in arrest. Transcriptional quiescence and stability of germline transcripts may aid in maintaining germline integrity and identity, supporting reproductive success of arrested larvae upon recovery.

**Figure 7:**
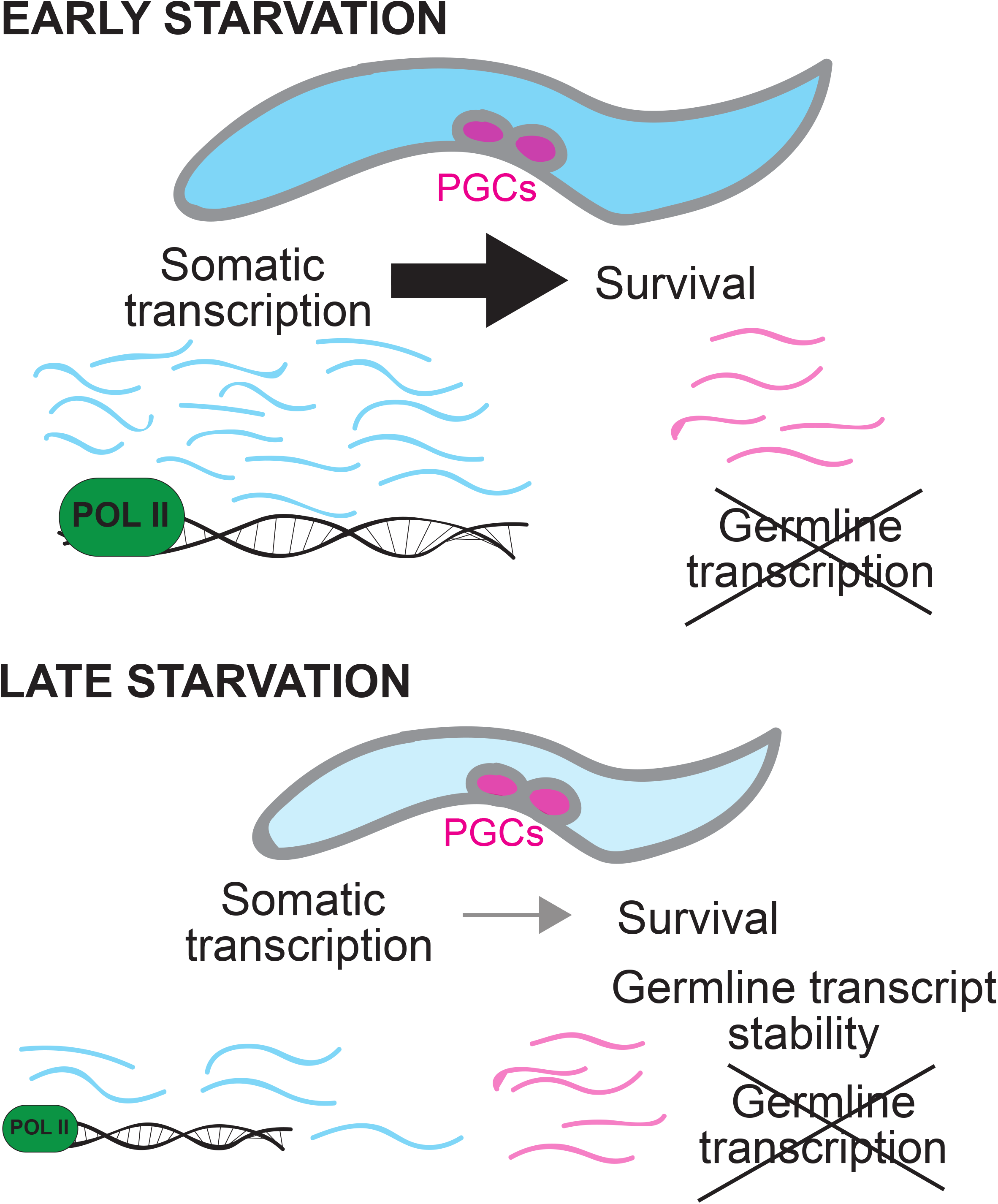
Somatic and germline gene regulation during starvation-induced developmental arrest. After hatching in the absence of food, L1-stage larvae arrest development and mount a transcriptional response in the soma that is required for survival. Ongoing somatic transcription maintains the initial response, but such late transcription has little impact on survival. In contrast, the germ line remains essentially transcriptionally quiescent throughout arrest. However, germline transcripts are particularly stable, and their relative expression levels increase while the abundance of other gene products decreases throughout the animal. These observations reveal alternative strategies for somatic and germline gene regulation during developmental arrest.

### Gene expression continues changing deep into starvation-induced developmental arrest

Gene expression analysis of starvation and developmental arrest typically focuses on a relatively early phase of the response (Wang and Kim 2003; Baugh *et al.* 2009; Sinha *et al.* 2012). Such work has revealed pervasive effects of nutrient availability on gene expression, but expression dynamics deep into cellular quiescence and developmental arrest have not been characterized. Our timeseries analysis reveals a rapid, widespread response within hours of hatching in the absence of food followed by a gradual but dramatic decline in the rate of transcriptome change. Nonetheless, gene expression levels continue changing throughout starvation, with a few thousand genes displaying apparent up-regulation in whole worms deep into starvation. Notably, ~84% of detected genes are differentially expressed over time during starvation, highlighting the vast physiological changes that take place.

Gene set enrichment analysis revealed differences in temporal expression patterns and regulation of genes with expression biased toward the soma or germ line. We used multiple published data sets that specify genes with expression in germ or somatic cells (Angeles-Albores *et al.* 2016; Lee *et al.* 2017; Angeles-Albores 2018; Packer *et al.* 2019) to associate tissues or cell types with temporal expression patterns. Surprisingly, many genes expressed in the germ line display peak expression levels deep into starvation. This was generally in contrast to genes expressed in the soma, whose expression tends to peak early. However, genes expressed in muscle are a notable exception in that they also tend to peak late. We also found that genes with a poised form of Pol II (docked Pol II) (Maxwell *et al.* 2014) are enriched among genes expressed in the germ line and genes whose expression peaks late. Again, this is in contrast to soma-expressed genes, which are instead associated with an active, elongating form of Pol II. These intriguing patterns of germline gene expression and regulation raise important questions. For example, why are genes with an inactive, docked form of Pol II increasing in expression late into starvation, and why are germline-expressed genes displaying this pattern in particular?

### Somatic transcription is required early in arrest to promote starvation survival

The dramatic and essentially immediate gene-expression response to starvation suggests a critical role of early transcription. Indeed, genes regulated by DAF-16/FoxO, DAF-18/PTEN, AMPK, LIN-35/Rb, and HLH-30/TFEB, each of which contributes to transcriptional regulation and is essential to starvation survival, are differentially expressed within hours of hatching without food. These results are consistent with a critical role of early transcription in mounting the starvation response, and DAF-16/FoxO nuclear localization dynamics support this interpretation (Weinkove *et al.* 2006; Mata-Cabana *et al.* 2020). To go beyond correlation, we used the AID system to conditionally degrade AMA-1/Pol II starting at different times in L1 arrest. Critically, transcription is required relatively early to support survival days later, with its requirement easing between 36 and 84 hr. The transcriptional inhibitor alpha-amanitin corroborated these findings. These results functionally demonstrate the importance of transcription in initially mounting the starvation response. mRNA-seq together with AMA-1/Pol II degradation revealed ongoing transcription of thousands of genes late in arrest, but expression of these genes peaks early as if their ongoing transcription reflects maintenance of the starvation response. In contrast to developmental gene-regulatory networks, which produce expression cascades, the lack of a distinct late starvation response suggests a relatively shallow regulatory network. Notably, degradation of AMA-1/Pol II with tissue-specific TIR1 transgenes revealed that somatic transcription promotes starvation survival but that germline transcription does not affect survival. These results are consistent with transcription deploying and actively maintaining the starvation response in somatic tissues while the germ line is transcriptionally quiescent and/or irrelevant to the survival of the animal (see below).

### The germ line remains transcriptionally quiescent throughout developmental arrest

Since it was a surprising observation, we focused on the apparent up-regulation of germline-expressed genes late in starvation. The germ line is largely transcriptionally quiescent throughout embryogenesis (Seydoux *et al.* 1996; Seydoux and Dunn 1997; Wang and Seydoux 2013), and bulk zygotic genome activation (ZGA) occurs in response to feeding in L1 larvae (Furuhashi *et al.* 2010; Butuci *et al.* 2015). DAF-18/PTEN and AMPK are both required to prevent PGCs from dividing and to support L1 starvation survival (Fukuyama *et al.* 2006; Fukuyama *et al.* 2012). DAF-18 inhibits ZGA in germ cells (Fry *et al.* 2020), and AMPK mutants have abnormally high levels of chromatin marks for transcriptional activation in PGCs of starved L1 larvae (Demoinet *et al.* 2017). In addition, *lin-35/Rb* mutants die rapidly during L1 arrest and express germline genes in the soma (Wang *et al.* 2005; Petrella *et al.* 2011; Wu *et al.* 2012; Cui *et al.* 2013). Does germline ZGA occur deep into L1 arrest, or are germline genes ectopically expressed in the soma, as if transcriptional repression eventually fails in wild type as it does in these mutants? Ablation of the germ line extends adult lifespan (Hsin and Kenyon 1999), and many common pathways regulate starvation resistance and aging (Baugh 2013), supporting the possibility that blocking transcription of germline genes could actually improve starvation survival. Despite these possibilities, our data shows that degrading germline Pol II during L1 arrest does not affect gene expression, indicating that PGCs remain transcriptionally quiescent throughout L1 arrest. Furthermore, degradation of germline Pol II does not affect starvation survival, and degradation of somatic Pol II does not increase survival. These results argue against the possibility of germline gene expression late in starvation, in the germ line or soma, compromising survival. PGCs are competent for transcription, as evidenced by feeding immediately stimulating germline ZGA and our observation that Pol II is required during L1 arrest for germline development upon feeding. Thus, L1 PGCs exhibit robust regulation to ensure transcriptional quiescence deep into arrest, and apparent up-regulation of germline genes and death in wild-type worms is not merely a recapitulation of the mis-regulation seen in certain mutants.

This work reveals that the PGCs in starved post-embryonic animals behave like germline blastomeres during embryogenesis by maintaining global repression of transcription while stabilizing transcripts that define germ-cell identity. PIE-1 is involved in both transcription inhibition and transcript stabilization in germline precursors (Tenenhaus *et al.* 2001; Wang and Seydoux 2013), making its potential involvement during L1 starvation enticing, but because it is eventually degraded in embryonic PGCs (Mello *et al.* 1996), it is likely that other factors are involved. Transcriptional repression in L1 PGCs during starvation appears to require DAF-18/PTEN and AMPK, with AMPK antagonizing the COMPASS complex (Demoinet *et al.* 2017; Fry *et al.* 2020), but additional molecular mechanisms remain to be determined. Transcriptional quiescence of germline cells, and possibly other stem cells, during starvation is likely conserved among metazoa.

### Transcript stability maintains germline gene expression deep into starvation

mRNA-seq measures steady-state mRNA expression levels, and differential expression may be due to differences in transcription, transcript stability, or both. Many studies assume that up-regulation is due to transcription, which is a reasonable assumption if the expression levels of most genes are not changing. However, it is difficult to know at the outset of an experiment whether this is the case, and notable exceptions exist (Radonjic *et al.* 2005; Lin *et al.* 2012). We found that germline-expressed genes increase relative expression late in arrest regardless of whether Pol II is present. This suggests that apparent up-regulation of germline genes is driven by transcript stability rather than transcription. Notably, expression of these genes is enriched in germ cells, but it is not exclusive to germ cells, so some ‘germline genes’ may be stabilized in the soma. In addition to using the AID system to conditionally eliminate transcription in the soma or germ line, we used 92 synthetic transcripts at known concentrations as spike-in standards with mRNA-seq. This approach enabled us to infer the relative fraction of total RNA represented by poly-adenylated mRNA in each sample, and it allowed us to convert read counts to absolute estimates of transcript abundance per worm. Total RNA per worm declines deep into starvation, and the mRNA-to-total RNA ratio remains approximately constant, suggesting germline genes remain relatively stable as the overall transcriptome declines, accounting for their apparent up-regulation in whole worms. Notably, we also found that protein levels decrease during L1 arrest, and we previously reported that arrested L1s shrink (Hibshman 2017), consistent with autophagy and other processes supporting survival but contributing to somatic collapse as starved larvae approach death. Together these approaches demonstrate that the amount of mRNA per worm decreases deep into starvation and that germline-enriched transcripts are generally more stable than those enriched in the soma.

Blocking transcription genetically or pharmacologically is a common way to measure the stability of transcripts in yeast (Herruer *et al.* 1988; Wang *et al.* 2002), and interpretation of relative up-regulation as increased stability when transcription is inhibited has precedent (Grigull *et al.* 2004). This interpretation is also corroborated by the association of germline genes with docked, but not active, Pol II. Eukaryotic mRNA half-lives can range from minutes to days (Lugowski *et al.* 2018), and mRNA half-lives can be increased in response to stress (Hollams *et al.* 2002). In our study, adequate mRNA was available for generating RNA-sequencing libraries with a standard protocol for at least four days after Pol II degradation, suggesting exceptional transcript stability in a starved, developmentally arrested state. Some studies have linked transcript degradation to translation (Radhakrishnan and Green 2016), and it is possible that transcripts important for germ cell identity and/or initiation of PGC proliferation (such as maternal germline transcripts) tend not to be translated during starvation allowing them to more easily escape degradation than transcripts important for survival.

### Alternative somatic and germline gene-regulatory strategies during developmental arrest

Our results demonstrate differential regulation of somatic and germline genes deep into starvation. These differences mirror consequences of extended L1 arrest on recovery, growth, and fecundity. In particular, larvae exhibit reduced growth rate and produce smaller broods following extended L1 arrest (Jobson *et al.* 2015). They are also more likely to develop gonad abnormalities including hyperproliferative germline tumors, which largely account for the reduction in brood size (Jordan *et al.* 2019). Single-worm expression analysis also suggests a mismatch between somatic and germline developmental rates following L1 starvation (Perez *et al.* 2017). The mechanistic basis of this mismatch is unclear, but increased maternal provisioning of vitellogenin lipoprotein and decreased larval insulin-like signaling decrease the mismatch, buffering development from L1 starvation (Perez *et al.* 2017; Jordan *et al.* 2019; Olmedo *et al.* 2020).

L1 arrest provides a unique opportunity to study trade-offs and mechanisms of somatic and germline maintenance in an enduring quiescent organismal state. Unlike other aging models, resources are finite in an essentially closed system. Stabilization of germline transcripts, and presumably other gene products, while the transcriptome, proteome, and body size decline, is reminiscent of the disposable soma theory of aging. This theory posits trade-offs between somatic maintenance and reproduction, such that the soma collapses in adults as reproduction is prioritized (Van den Heuvel *et al.* 2016). Dauer larvae and arrested L1s are pillars of prioritizing somatic maintenance, as they postpone reproduction and survive for weeks or months without developing (Baugh and Hu 2020), and they do so with minimal effects on lifespan upon recovery (Klass and Hirsh 1976; Johnson *et al.* 1984). However, markers of aging accumulate during L1 arrest, including protein aggregates, DNA damage, reactive oxygen species, reduced motility, body shrinkage, and reduced mitochondrial function, and many of these aging markers are actively reversed upon recovery (Roux *et al.* 2016b; Hibshman *et al.* 2018), causing a lag in the initiation of post-embryonic development (Olmedo *et al.* 2020). Here we show that the germ line is maintained in a transcriptionally inactive state and that germline transcripts display exceptional stability as the soma collapses. We believe that such germline regulation provides passive resistance to nutrient stress, preserving germline integrity and identity. From an evolutionary perspective, this makes sense – maintenance of the germline is the only way to ensure subsequent generations. Furthermore, it is possible that by remaining so quiescent, germ cells dodge aging during arrest, thereby requiring less time to recover and initiate proliferation than somatic cells upon recovery, thus accounting for the mismatch in somatic and germline developmental rates resulting from L1 arrest. However, this raises the question of why the soma has an active response to starvation, rather than adopting a deeper state of quiescence by entering a cryptobiotic state. Such a state could enable extreme endurance but it would also result in behavioral quiescence, eliminating the animal’s ability to forage, feed, and ultimately recover from starvation. Thus, the different responses to starvation displayed by the soma and germ line reflect different demands placed on each in order to support organismal fitness.

## STAR METHODS

### Resource Availability

#### Lead contact

Further information and requests for resources and reagents should be directed to and will be fulfilled by the Lead Contact, Ryan Baugh.

#### Materials availability

Strains generated in this study are available upon request.

#### Data and code availability

mRNA-seq datasets generated in this study are available at NCBI GEO at GSE173657.

### Experimental model and subject details

#### Strains used

N2 was obtained from the Sternberg collection at the California Institute of Technology, originally from the CGC in 1987. Strains obtained from the CGC are:

CA1200 ieSi57 [*eft-3p*::TIR1::mRuby::*unc-54* 3’UTR + Cbr-*unc-119*(+)] II – referred to as *Peft-3::TIR1* CA1202 ieSi57 II; ieSi58 [*eft-3p*::degron::GFP*::unc-54* 3’UTR + Cbr-*unc-119*(+)] IV – referred to as *Peft-3::TIR1; Peft-3::AID:GFP*

CA1352 ieSi64 [*gld-1p*::TIR1::mRuby::*gld-1* 3’UTR + Cbr-*unc-119*(+)] II – referred to as *Pgld-1::TIR1*

Newly generated strains from this study are:

PHX1513 *ama-1(syb1513)* – referred to as *ama-1::AID*

LRB387 ieSi64 II; *ama-1(syb1513)* IV – referred to as *Pgld-1::TIR1; ama-1::AID*

LRB389 ieSi57 II; *ama-1(syb1513)* IV – referred to as *Peft-3::TIR1; ama-1::AID*

### Method details

#### Sample preparation and collection for mRNA-seq time series

N2 was maintained well-fed on OP50 at 20°C for at least three generations. For each biological replicate, seven to ten gravid adults were picked onto approximately 20 large (10 cm) NGM plates with OP50 3.5 days prior to hypochlorite treatment (bleach). Large plates were then hypochlorite treated as described previously (Hibshman *et al.* 2021) to obtain over 200,000 embryos per biological replicate. Embryos were resuspended in S-basal at a density of 1 embryo/μl in an Erlenmeyer flask at 20°C in a shaker at 180 rpm. Beginning at 10 hours after hypochlorite treatment, hatching efficiency was scored. Time points used in the time series are 12 hours offset from hypochlorite treatment and signify when worms have begun hatching and are thus undergoing L1 arrest. L1 larvae hatch approximately 12 hours after hypochlorite treatment, and this is considered 0 hr. Scoring continued every two hours until the curve appeared to level off at 18 hours after hypochlorite treatment, or six hours of L1 arrest (Fig 1B). At least 10,000 L1s were collected per time point. For days 8 and 12, the number of L1s collected was doubled and quadrupled, respectively, to account for lethality at later time points. To collect samples, L1s in S-basal were first spun down at 3,000 rpm, and S-basal was aspirated off down to less than 100 μL. The pellet and residual S-basal was transferred to a 1.5 mL Eppendorf tube using a glass pipet. Samples were flash frozen in liquid nitrogen and stored at −80°C until RNA isolation.

#### RNA isolation and library preparation for mRNA-seq time series

RNA was extracted using TRIzol Reagent (Thermo-Fisher Scientific) using the manufacturer’s protocol with some exceptions. 100 μL of sand (Sigma-Aldrich) was included to aid homogenization. Sand was first prepared by washing two times in 1 M HCl, washed eight times in RNAse-free water (to a neutral pH), and baked to dry. Libraries were prepared for sequencing using the NEBNext Ultra RNA Library Prep Kit for Illumina (New England Biolabs, E7530) with 100 ng of starting RNA per library and 14 cycles of PCR. Libraries were sequenced using Illumina HiSeq 4000. Four biological replicates were sequenced per time point. Three replicates consisted of samples collected from the same culture (12 time points collected from a single culture). One replicate consisted of samples from multiple original cultures.

#### Differential expression analysis for mRNA-seq time series

Single-end 50 bp reads were mapped with with bowtie (Langmead 2010) using the following command: bowtie --best --chunkmbs 256 -k 1 -m 2 -S -p 2. Average mapping efficiency was 81.5% with a standard deviation of 2.0%. HTseq was used to count reads mapping to genes (Anders *et al.* 2015) using version WS210 of the *C. elegans* genome obtained from Maxwell et al. (Maxwell *et al.* 2012). Count tables were used to detect differential expression in R using edgeR (Robinson *et al.* 2010). Prior to differential expression analysis, count tables were filtered first to include only genes with counts per million (CPM) greater than one across four libraries. This resulted in 16,699 genes for further analysis. The edgeR glm functionality was used to fit a GLM, and a likelihood ratio test was performed for each gene; this ANOVA-like test was used to detect any differences across the twelve time points. PCA was done on log2 mean-normalized CPM values for the 16,699 detected genes (Fig 1D). For rate-of-change analysis, the Euclidean distance between all replicates for adjacent time points (using the same input as for PCA) was calculated and divided by the duration of time between the time points (Fig 1E).

#### Clustering and heatmap of time series

6,027 genes with FDR < 1 x10^−30^ were included in cluster analysis. Using the twelve average CPM values over time for each gene, the pairwise Pearson correlation coefficient was calculated for each gene compared to every other gene. One minus the Pearson correlation coefficient was calculated, and these values were assembled into a square matrix to use as input for clustering. Clustering was performed in Matlab, using a published algorithm (Heyer *et al.* 1999; Baugh *et al.* 2003). As parameters for clustering, 0.2 was used as the maximum distance for two genes to still be called in the same cluster, where the “distance” is again defined as one minus the Pearson correlation coefficient. Thus, genes in the same cluster have at minimum a Pearson correlation coefficient of 0.8. This non-deterministic analysis resulted in 129 clusters, with approximately two-thirds of genes in the top 10 clusters and smaller clusters containing as few as two genes.

To plot genes within clusters, the Z-score for each gene over the 12 time points was calculated. The Z-score scales expression by the mean and standard deviation rather than abundance. Plots were generated for each cluster in R using the package ggplot2. To generate heatmaps, the R package pheatmap was used. 6,027 genes used for clustering were plotted in the heatmap. Genes within the same cluster were plotted next to each other without additional sorting. Clusters were first sorted by time point of highest average expression assessed by Z-score. Clusters with the same time point of highest expression were sorted by their ‘centroid’ time point. The centroid was calculated by transforming z-scores for each cluster so that the minimum was 0, calculating the sum across all 12 time points, and calculating the first time point at which the sum of the transformed Z-scores is less than half of the total sum. For clusters with the same maximum and centroid time point, clusters were sorted based on the value of the maximum Z-score. Clusters with peak expression at earlier time points were plotted before those at later time points. Among clusters with the same peak expression time point, clusters were ordered by the time point. For clusters with peak expression at the same time point, clusters were ordered based on the Z-score at the peak. Clusters were ordered from highest to lowest for peaks in the first six time points, and from lowest to highest for peaks in the last six time points.

#### Gene enrichment analysis for gene groups of interest

Gene groups were either obtained from datasets in previous publications (including those used in Fig 2C-F and Fig 3) or generated as part of this study (Fig 6). For RNAseq datasets, genes down-regulated in a transcriptional regulator mutant relative to wild-type were considered positive targets, and genes up-regulated were considered negative targets. Wormbase IDs were used for clustered genes and gene groups of interest. If Wormbase IDs were not included in the original publication’s dataset, then Simplemine was used to convert gene identifiers. Gene groups were filtered to include only genes that were considered expressed in the time series dataset. For each cluster, the overlap between the filtered gene group and the cluster’s genes was determined, and a hypergeometric p-value was calculated. For plots including the mean Z-score over time with 99% confidence intervals (*i.e*., Fig 2), the stat_summary function in ggplot2 was used, and bootstrapping (mean_cl_boot) was the method for determining the confidence interval surrounding the mean. For plots including individual trajectories for all genes in the filtered gene group included in clustering (*i.e.* Figs. 3 and 6), lines were color-coded if the gene was part of an enriched cluster. Clusters were considered enriched if p < 0.01 or p < 0.05, as indicated in the figure legend. A more stringent p-value was used with larger gene groups. Genes in non-enriched clusters were still plotted, but with less opacity to enable visualization of all gene trajectories.

#### Design and generation of ama-1 AID strains

We designed PHX1513 *ama-1(syb1513*) to have an insertion prior to the stop codon at the endogenous *ama-1* locus. This insertion contained the degron sequence, linkers, and 3x-FLAG tag, in line with designs used to create other AID strains in *C. elegans* (Zhang et al. 2015; Kasimatis et al. 2018). The sequence of the insertion used was: GGATCCGGAGGTGGCGGGATGCCTAAAGATCCAGCCAAACCTCCGGCCAAGGCACAAGTTGTGGGATGGCCACCGGTGAGATCATACCGGAAGAACGTGATGGTTTCCTGCCAAAAATCAAGCGGTGGCCCGGAGGCG GCGGCGTTCGTGAAGGAGAATCTGTACTTTCAATCCGGAAAGGACTACAAAGACCATGACGGTGATTATAAAGATCATGATATCGATTACAAGGATGACGATGACAAGTAA

SunyBiotech generated the strain PHX1513 and validated it with Sanger sequencing. We also validated the strain upon receipt using Sanger sequencing. To generate the functional AID strains LRB387 and LRB389, PHX1513 was backcrossed twice to N2 and then crossed to strains expressing TIR1 in either the soma (CA1200) or germ line (CA1352).

#### Auxin preparation and treatment

A 400 mM stock solution of indole-3-acetic acid (IAA), or auxin, was prepared in ethanol and stored at - 20°C. For experiments using a 0.1 mM auxin dose, a 100 mM stock in ethanol was used, and a corresponding amount of ethanol was used (0.1%) for the control. For experiments using a 1 mM auxin dose, the 400 mM stock was used, and a corresponding amount of ethanol (0.25%) was used for the control.

#### Starvation survival assays

Starvation survival in Fig 1C was assayed by indirect scoring (Baugh and Hu 2020), which considers worms alive if they are able to recover upon feeding, as done previously (Webster *et al.* 2018). 100 μL of each 8-day and 12-day sample used for RNA-seq was aliquoted around the edge of a 5 cm NGM plate with a spot of OP50 in the center, and the number of individuals plated was counted. Survivors were scored two days later, and proportion alive was calculated.

For Fig 4F-G, direct scoring in arrested L1s was used to assay survival because worms with degraded AMA-1 in the soma could not recover. To prepare plates to bleach, ~10 L4s from *Peft-3::TIR1*, *Pgld-1::TIR1*, and *Pgld-1::TIR1;ama-1::AID* were picked onto approximately four 10 cm plates with OP50. ~15 adults were picked from *Peft-3::TIR1;ama-1::AID* onto at least seven 10 cm plates with OP50 to account for slower growth and fewer progeny from this strain. Four days later, gravid adults from each genotype were hypochlorite treated from unstarved plates. S-basal with 0.1% ethanol was used throughout washing and hypochlorite treatment. For each condition (genotype and auxin or ethanol addition at each time point), a 5 mL culture of S-basal with 0.1% ethanol was set up in a 25 mL Erlenmeyer flask at 1 embryo/μL (5,000 embryos total) and stored in a 20°C shaker moving at 180 rpm. At the indicated time point, 0.1 mM auxin was added for +auxin conditions, and 0.1% ethanol was added for ethanol conditions. This ensured an equal amount of ethanol (final concentration of 0.2%) across all conditions. After 12 days of L1 arrest, at least 100 μL was sampled from the culture and pipetted on to a glass depression slide. Worms were scored as dead if they were rods or lacked muscle tone and had a granular appearance without apparent movement. Proportion alive was calculated as the number of live worms divided by the sum of alive and dead worms.

For starvation survival with alpha-amanitin (Supplementary Fig 4C), direct scoring was also used. N2 worms were hypochlorite treated and embryos were resuspended in S-basal at 1 embryo/μL in Erlenmeyer flasks and placed in a shaker, as was done with the *ama-1::AID* strains. 12 hours after hatch (24 hours after hypochlorite treatment), 25 ug/mL alpha-amanitin was added to cultures. Survival was scored on days 2 and 8, with day 8 as an earlier last time point than with AMA-1 AID because ethanol was not added to the cultures.

#### Starvation recovery

For Fig 4C, strains were scaled up and hypochlorite treated as described in ‘Starvation survival assays,’ but without ethanol added to S-basal. Embryos were resuspended at 1 embryo/μL in 5 mL of S-basal in a 16 mm glass test tube and stored on a roller drum at 20°C. After 23 hours (11 hours of L1 arrest), 1 mM auxin or 0.25% ethanol was added to each test tube. After one hour, worms were transferred to 15 mL conical tubes and spun down at 3,000 rpm, and excess S-basal was aspirated off. S-basal was used to wash worms three additional times, then they were plated on 10 cm NGM plates with OP50. After 48 hours of growth, worms from each condition were washed off of plates with S-basal, spun down, and transferred to an unseeded 10 cm NGM plate for imaging. Images were taken on a ZeissDiscovery.V20 stereomicroscope and analyzed using the WormSizer plugin for FIJI (Moore *et al.* 2013).

#### Hatching efficiency

5 cm NGM plates with OP50 were prepared with different doses of auxin (1, 0.1, 0.01, 0.001, 0.0001 mM) by pipetting the appropriate amount of auxin and ethanol onto an already-seeded plate immediately before use. A cell spreader was used to quickly and lightly (to minimize interfering with the lawn) spread the auxin on the plate. Doses of auxin were prepared from the 400 mM auxin stock, and lower doses were diluted in ethanol so that the same amount of ethanol was used for each dose. Embryos resuspended in S-basal following hypochlorite treatment were pipetted onto the plates prepared for each dose. After 24 hours, the number of embryos and L1s were scored on each plate. 100-300 individual animals were scored for each replicate and condition.

#### Gonad cell scoring upon recovery

Approximately 10 *Peft-3::TIR1* and *Peft-3::TIR1; ama-1::AID* L4s were picked onto each of 3-4 10 cm NGM plates seeded with OP50. 4 days later, strains were hypochlorite treated as described for ‘Starvation survival assays’, with S-basal with 0.1% ethanol included throughout. 5 mL cultures were set up with embryos resuspended at 1 embryo/μL in 16 mm glass test tubes. Immediately after cultures were set up (within approximately one hour post-bleach), 0.1 mM auxin or 0.1% ethanol was added to cultures for the 48 hours exposure conditions. Test tubes were then put on a rotating roller drum at 20°C. At 24 hours post-bleach (approximately 12 hours of L1 arrest), 0.1 mM auxin or 0.1% ethanol was added to cultures for the 24-hour exposure conditions. At 48 hours post-bleach, all six cultures (both 24-hour and 48-hour exposures) were transferred to 15 mL conical tubes, spun down at 3,000 rpm, and excess S-basal was aspirated off down to < 100 μL containing the arrested L1s. Worms were washed six times with 10 mL S-basal. After all washes, worms were transferred to 10 cm NGM plates seeded with OP50. After 24 hours of growth on plates, 4% agar slides were prepared for viewing on a compound microscope. 2 μL of levamisole was pipetted onto each slide, individual worms were picked from plates into the levamisole, and a cover slip was placed. Individual cells in the gonad were then counted using an AxioImager compound microscope (Zeiss) at 1000x total magnification. DIC images were taken using Zen software (Zeiss), and cropped in Adobe Illustrator.

#### Sample preparation and collection for mRNA-seq on AMA-1 AID samples

All strains were maintained well-fed on OP50 at 20°C for several generations. For *Pgld-1;:TIR1; ama-1::AID*, *Peft-3::TIR1*, and *Pgld-1::TIR1*, approximately 10 L4s were picked onto five 10 cm NGM plates with OP50 for each biological replicate. For *Peft-3::TIR1; ama-1::AID*, which grows slower, approximately 15 gravid adults were picked onto seven 10 cm NGM plates with OP50 for each biological replicate. After 4 days, plates from all four strains were hypochlorite treated in parallel. For each strain, three 50 mL Erlenmeyer flasks were used, each with 10,000 embryos resuspended at 1/μL in 10 mL. Flasks were put in a shaker at 180 rpm at 20°C, like was done for the time series samples. After 36 hours of L1 arrest (48 hours after bleach), the 36-hour samples were collected by spinning down the 10 mL culture for each strain at 3,000 rpm, aspirating off excess S-basal, and then transferring the worms in ~100 μL to a 1.5 mL Eppendorf tube. The volume of the sample was further reduced by spinning again at 3,000 rpm and pipetting off excess liquid to bring the final volume down to approximately 10 μL. This small volume facilitates micro-scale RNA preparation. Samples were flash frozen in liquid nitrogen and stored at −80°C until RNA isolation. At 36 hours, 100 μM auxin or ethanol was added to each of the remaining flasks. After 132 hours of L1 arrest, auxin and ethanol samples were collected as described for the 36-hour samples.

#### RNA isolation and library preparation for AMA-1 AID samples

RNA was isolated using TRIzol Reagent (Invitrogen# 15596026) following the manufacturer’s instructions with some exceptions noted below. The procedure was scaled down linearly, using 100 μL Trizol. 5 ug linear polyacrylamide (Sigma# 56575) was included as a neutral carrier for RNA precipitation. RNA was eluted in nuclease-free water and incubated at 55ºC for approximately four minutes to resuspend the pellet. 25 ng of total RNA was used for each library preparation. ThermoFisher ERCC Spike-In Mix (Thermo #4456740) was added based on total RNA per manufacturer’s instructions. NEBNext Poly(A) mRNA Isolation Module (New England Biolabs# E7490) and NEBNext Ultra II RNA Library Prep kit (New England Biolabs# E7770) was used to perform poly-A selection and prepare libraries for sequencing respectively. The final libraries were enriched with 14 or 15 cycles of PCR, depending on batch. Libraries were then sequenced using Illumina NovaSeq 6000 to obtain 50 bp paired-end reads.

#### Differential expression analysis of AMA-1 AID RNA-seq data

Paired-end reads were mapped with bowtie using the following command: bowtie -I 0 -X 500 -- chunkmbs 256 -k 1 -m 2 -S -p 2 using the WS273 version of the genome, with the sequences from the ERCC spike-ins appended to determine reads mapping to the spike-ins. Counts were determined as described for the time series analysis. Count data was imported into R, then filtered to include only protein-coding genes. To determine expressed genes, count data was further filtered to include only genes with counts > 10 in at least four samples. The RUVseq package (Risso *et al.* 2014) was used to determine factors of unwanted variation prior to differential expression analysis. First, betweenLaneNormalization was used for upper-quartile normalization (which = “upper”). Next, RUVg was used with k = 1 to improve normalization by normalizing based on the spike-ins. edgeR was then used for differential expression analysis, incorporating the factors of unwanted variation from RUVseq. glmFit and glmLRT commands (which fit a GLM and execute a likelihood ratio test in edgeR) were used to determine significant differences for pairs of conditions of interest.

#### Normalization with external spike-in standards

Counts mapping to spike-ins were plotted against the known concentrations of spike-ins in Mix 1 added to total RNA. A linear model was fit in R between the log_10_-normalized counts and log_10_-normalized concentration of each spike-in. Based on the slope and intercept values from the regression fit, all counts for non-spike-in genes were converted to units of attomoles per μL (the spike-in units). To convert to units of attomoles of transcript per worm, the normalized count data was then multiplied by the volume (in μL) of spike-in and divided by the average number of worms needed to acquire the amount of total RNA used for each condition.

#### Lysate preparation for Western blot

Worm samples were collected as described for the mRNA-seq samples, with minor exceptions. Only the *Peft-3::TIR1; ama-1::AID* genetic background was used, and one of three biological replicates was collected with 25,000 instead of 10,000 worms for each condition (though density was held constant). Samples were stored at −80°C. The frozen pellet was rapidly freeze-thawed 3 times, cycling between liquid nitrogen and a 45°C water bath. Laemmli buffer (Sigma# S3401) was added to the samples to bring the final concentration to 1X. The samples were then incubated at 95°C for 10 minutes, frozen in dry ice for 15 minutes, and incubated at 95°C again for another 10 minutes. Debris was pelleted by centrifuging the samples at 14,000 rpm for 5 minutes, and the lysate was transferred to a new tube. Total protein was quantified using the Pierce 660nm Protein Assay kit (Thermo# PI22662) following the manufacturer’s instructions.

#### Western blot

Two μg of total protein per sample was loaded onto the NuPAGE 4%-12% Bis-Tris gel (Invitrogen# NP0321), along with 10 μl of Spectra Multicolor Protein Ladder (Thermo# 26634). Proteins were resolved at 200 V for 50 min in MOPS SDS running buffer. Following resolution, proteins were transferred to PVDF membrane (Invitrogen# LC2005) at 100 V for 60 min. The membrane was blocked in non-fat milk for 1 hour to reduce non-specific binding. The blot was then probed with HRP conjugated anti-FLAG antibody (Sigma# A8592) at 1:2000 through overnight incubation at 4°C. The membrane was then washed multiple times with TBS buffer, supplemented with 0.1% Tween-20, to wash off excess unbound antibodies. SuperSignal West Femto Substrate (Thermo# 34095) was used to develop the blot, and it was imaged in the iBright FL1500 Imaging system (Thermo Fisher). A second NuPAGE gel, run with the same samples, was stained with SYPRO Ruby Protein Gel Stain (Invitrogen# S12001) following the manufacturer’s protocol and imaged to confirm equal loading.

### Quantification and statistical analyses

#### Statistical analysis

Statistics were performed in R, and the statistical test used is indicated in figure legends. Linear mixed-effects models were used to control for the effect of biological replicate in cases in which there were multiple biological replicates and multiple observations originating from each replicate (*e.g.*, Figure 4E,G). Biological replicate was considered a random effect, and the variable of interest was considered a fixed effect. A t-test or one-way ANOVA was used in cases in which there was only one observation per replicate (*e.g.*, Figure 4B,H). The Kolmogorov-Smirnov test was used to determine if there were differences in distributions of a groups of genes across conditions (*e.g*., Fig 5D,F).

## ACKNOWLEDGEMENTS

We would like to thank E. Jane A. Hubbard for helpful discussions and Geraldine Seydoux and Chih-Yung Lee for sharing unpublished data analysis. This work was supported by the National Institutes of Health (R01GM117408, L.R.B.) Some strains were provided by the CGC, which is funded by NIH Office of Research Infrastructure Programs (P40 OD010440). AKW was supported by the NSF Graduate Research Fellowship Program.

## AUTHOR CONTRIBUTIONS

AKW and LRB designed experiments and wrote the paper. AKW and RC performed experiments. AKW analyzed data.

---

**Supplementary Figure 1:**
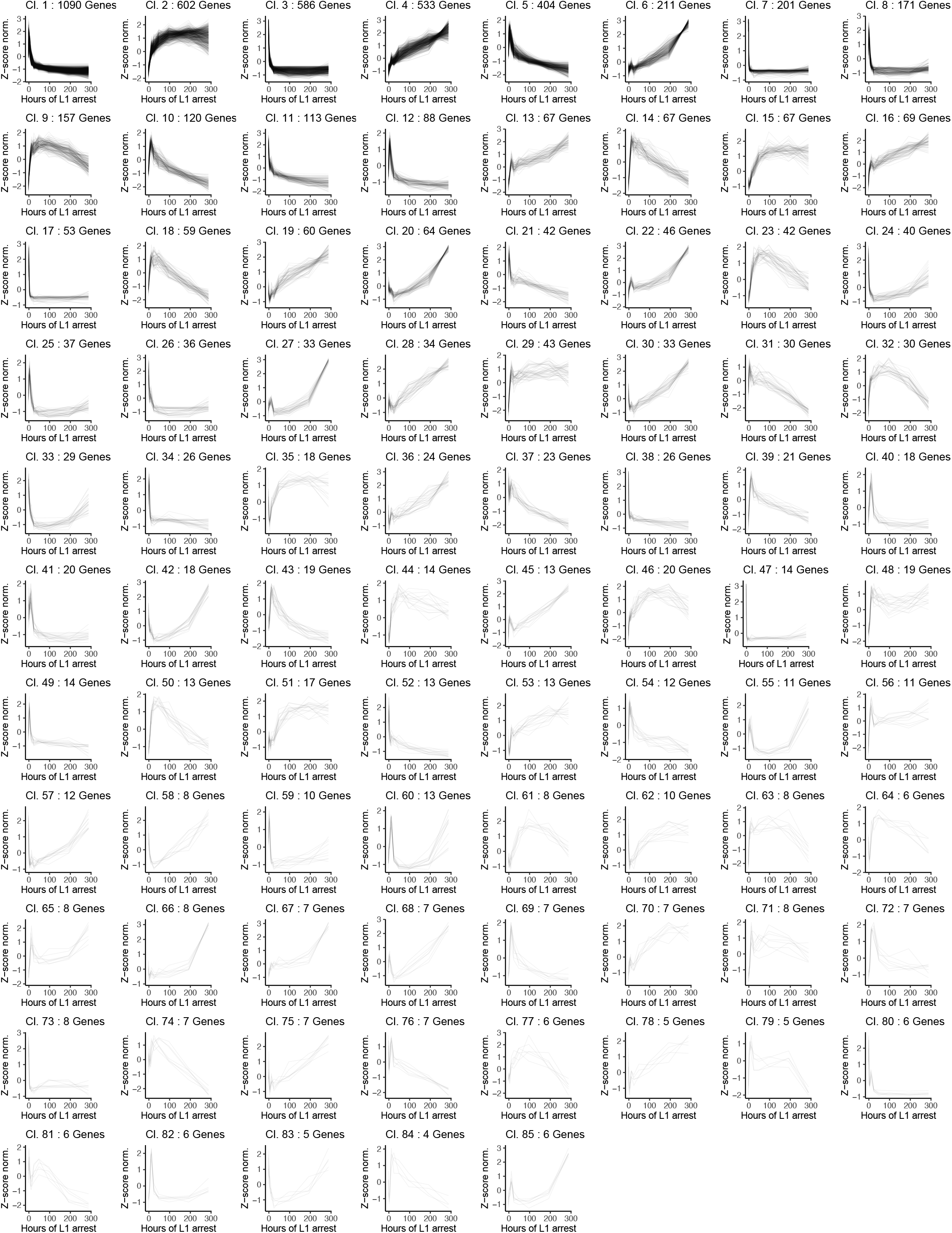
Cluster analysis reveals dominant temporal patterns of gene expression in whole starved worms. Z-scores over time for all genes included in Clusters 1-85 are shown, which includes all clusters with 5 or more genes. Clusters 86-129 are not shown, each of which has fewer than five genes.

**Supplementary Figure 2:**
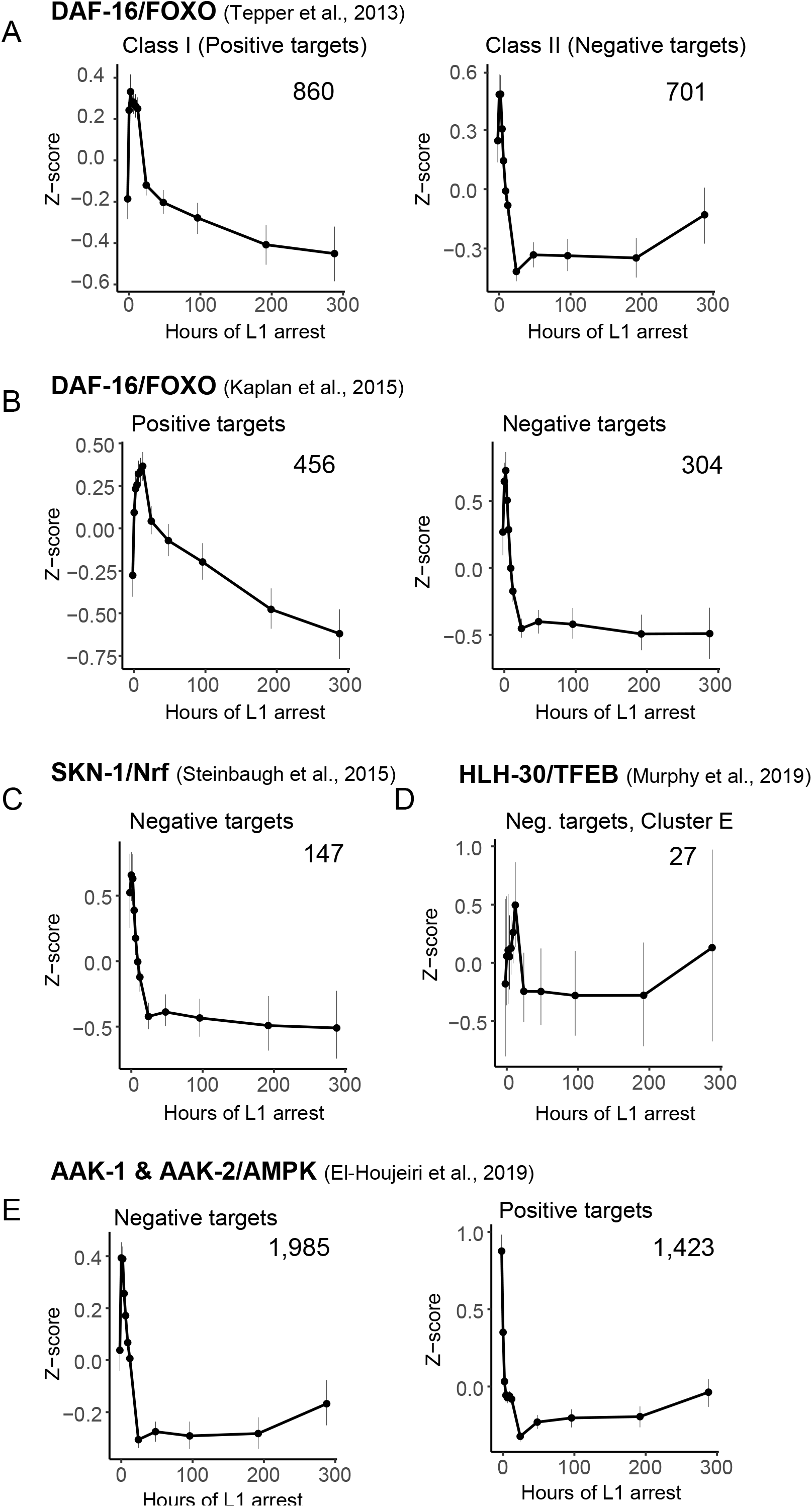
Temporal activity dynamics of additional transcriptional regulators. A-G. Average Z-scores over time for all genes included in clustering from the indicated regulator and cited dataset are plotted. Error bars show the 99% confidence interval surrounding the mean.

**Supplementary Figure 3:**
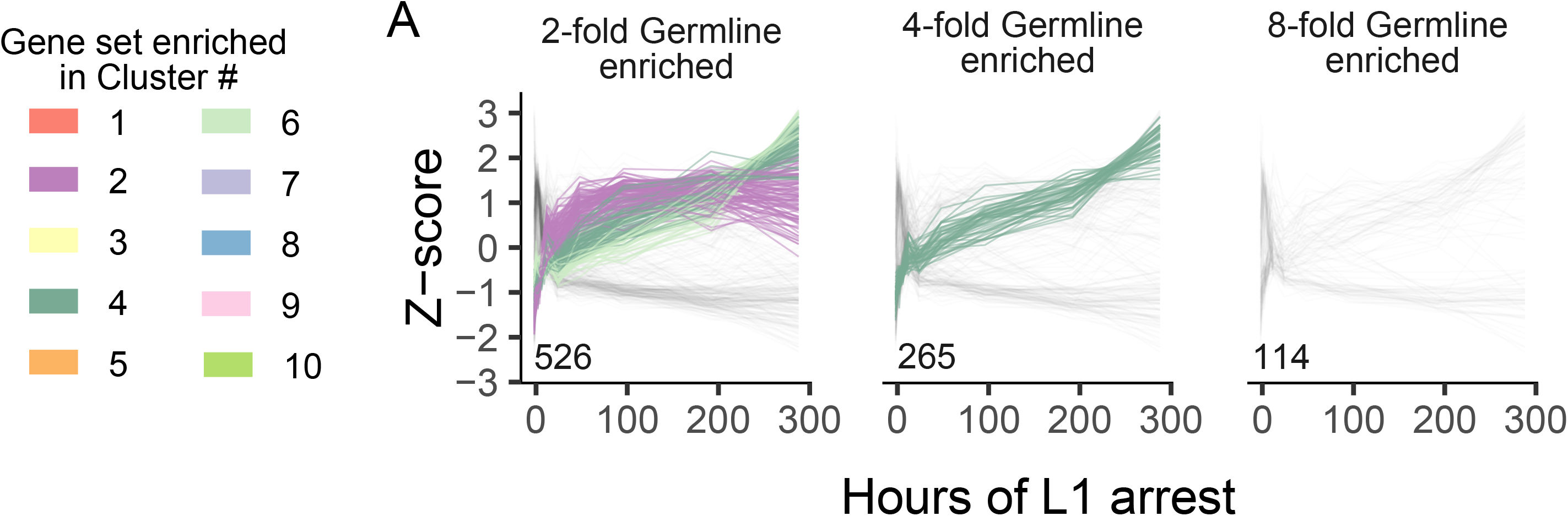
Relative increase in germline gene expression is robust to increased stringency in defining germline genes. A. Z-scores of all germline-enriched genes included in clustering are plotted over time. The number of genes plotted is indicated on the inset of each graph. Genes are color-coded by cluster if that cluster is enriched (hypergeometric p < 0.05) in the gene group.

**Supplementary Figure 4:**
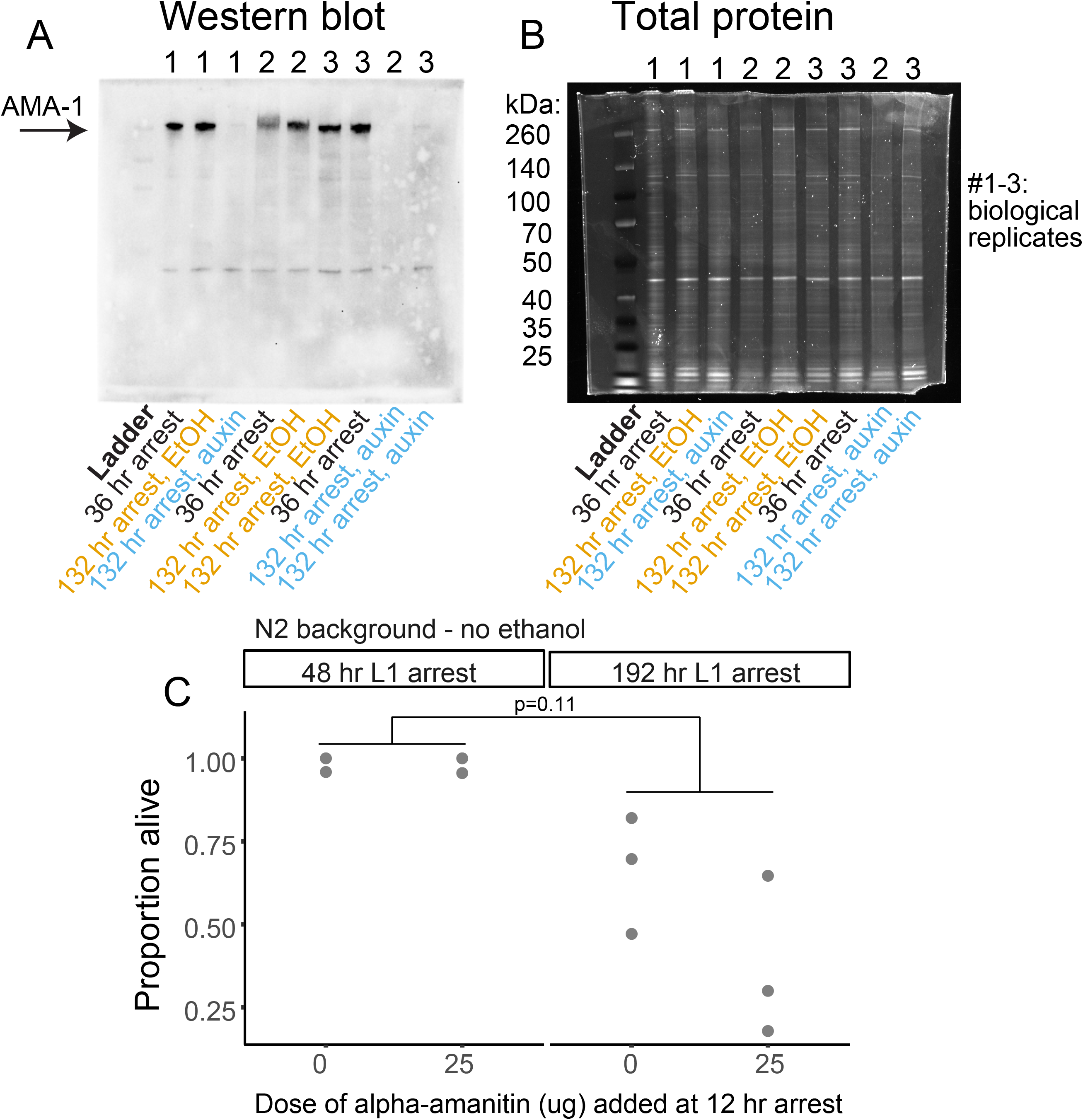
AMA-1 protein is degraded upon auxin addition during L1 arrest in *Peft-3::TIR1; ama-1::AID* background. A. Western blot probed with anti-FLAG to visualize AMA-1. Expected size of phosphorylated AMA-1 including the degron and 3xFLAG tag is ~218-258 kD. B. Total protein gel corresponding to Western blot in A. Equal amounts of total protein were loaded for all samples. A-B. The ladder and next three lanes are also shown in Figure 4D. C. N2 survival with and without addition of alpha-amanitin (Pol II inhibitor) addition. P-value calculated from linear mixed-effects model with interaction between time point and alpha-amanitin dose. Replicate was included as a random effect in the model.

**Supplementary Figure 5:**
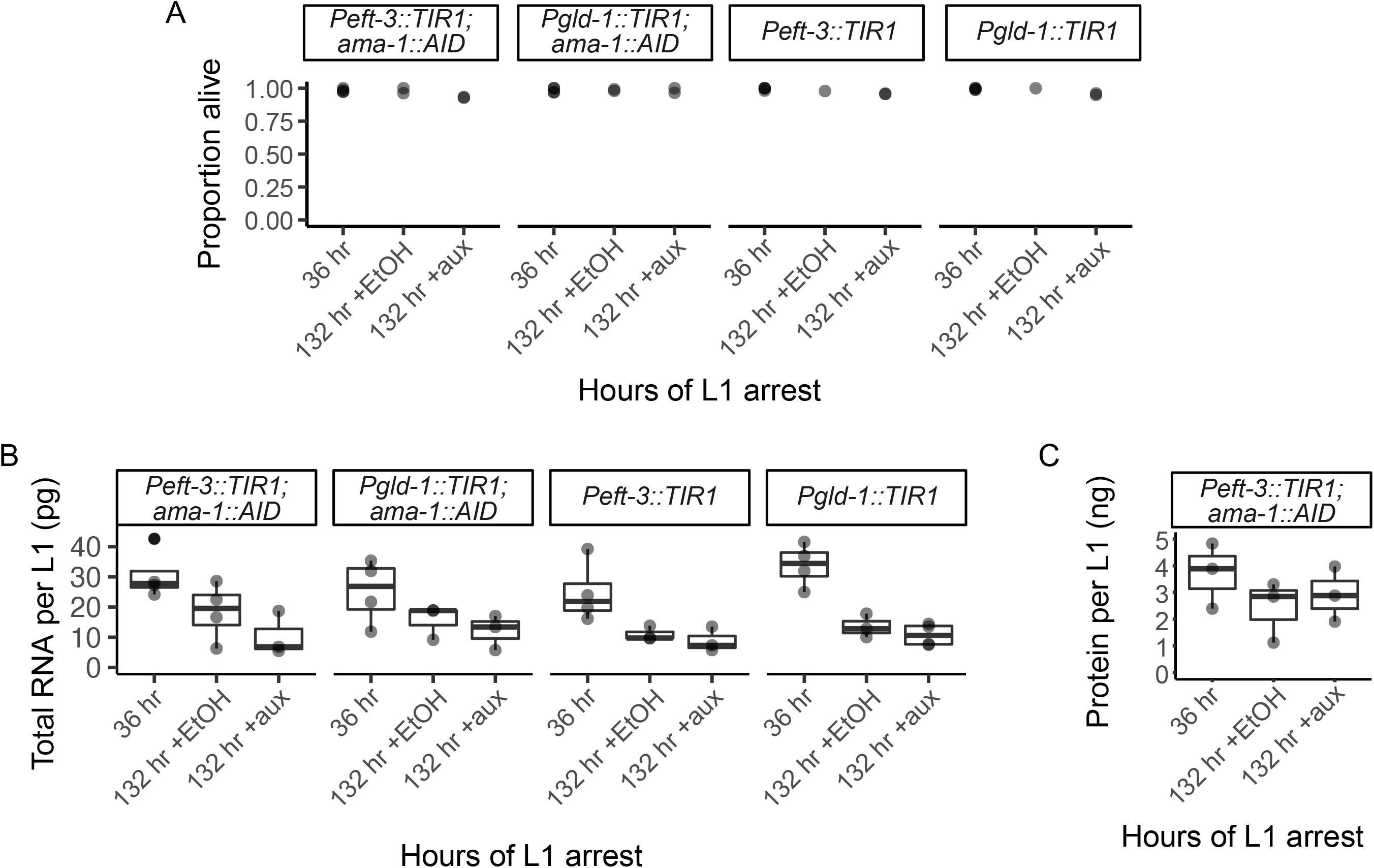
Survival, RNA content, and protein content in conditions used for *ama-1::AID* mRNA-seq. A. Proportion alive during L1 arrest at 36 hours, 132 hours with auxin, and 132 hours with ethanol. B. Total RNA content for all strains and conditions. One-way ANOVAs were performed within each time point across conditions and there were no differences. Data across strains was merged for Figure 6F. C. Total protein content across conditions in the *Peft-3::TIR1; ama-1::AID* background. There was no difference between 132 hours auxin and ethanol conditions in a t-test, so these were merged for Figure 6G.

**Supplementary Figure 6:**
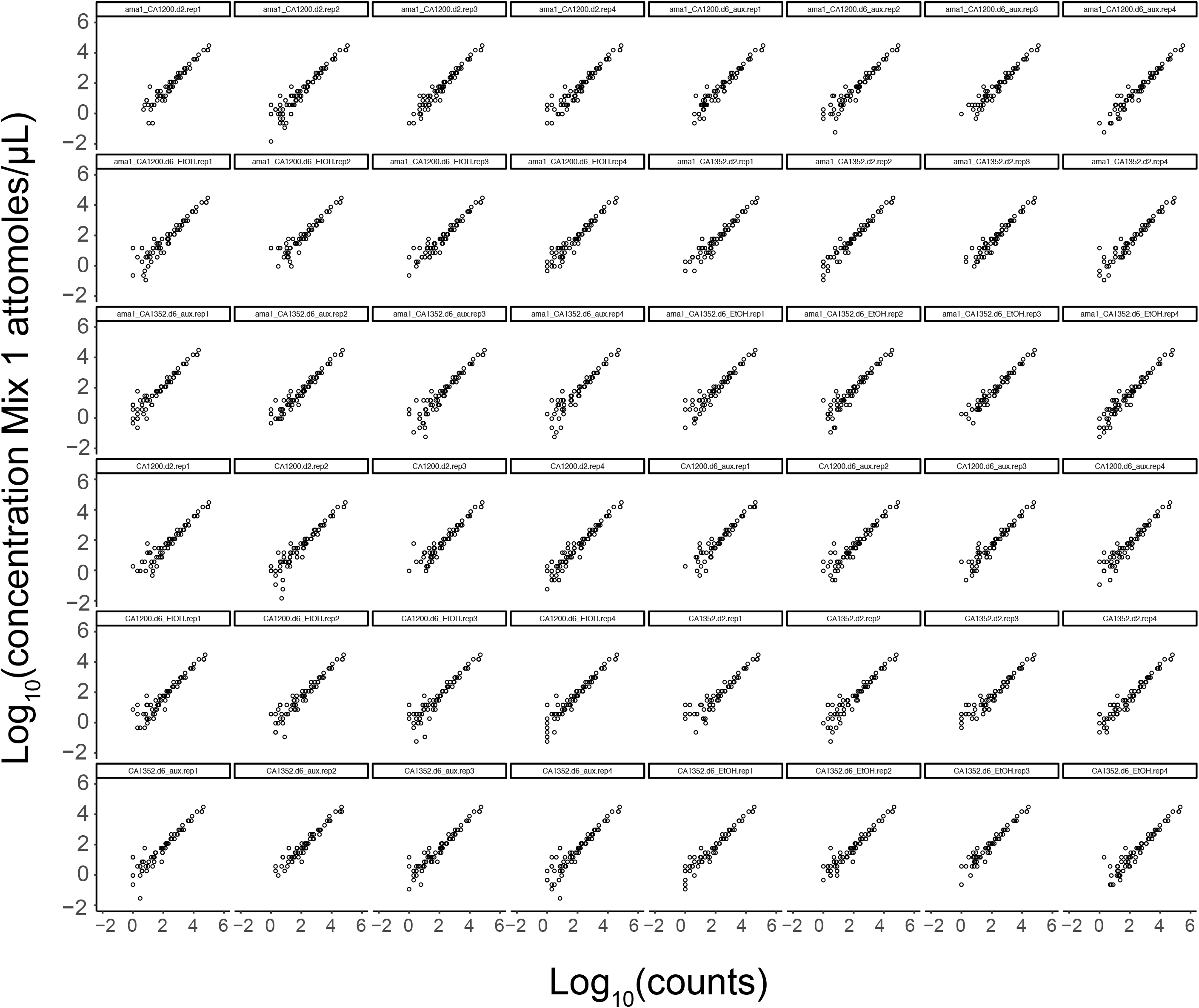
Known concentration of spike-ins vs. spike-in counts. mRNA-seq count data on the x-axis plotted against known concentration in attomoles per μL on the y-axis. A linear regression was fit to determine a normalization factor for each library for absolute normalization used for Figure 6H-I.

Supplementary File 1: Excel file with multiple sheets including count tables, CPMs, and differential expression output for both mRNA-seq experiments. Clusters, z-score normalized CPMs, and gene sets for enrichment analysis corresponding to the time series analysis are also included.

